# Evolution of multifunctionality through a pleiotropic substitution in the innate immune protein S100A9

**DOI:** 10.1101/865493

**Authors:** Joseph L. Harman, Andrea N. Loes, Gus D. Warren, Maureen C. Heaphy, Kirsten J. Lampi, Michael J. Harms

## Abstract

Multifunctional proteins are evolutionary puzzles: how do proteins evolve to satisfy multiple functional constraints? S100A9 is one such multifunctional protein. It potently amplifies inflammation via Toll-like receptor 4 and is antimicrobial as part of a heterocomplex with S100A8. These two functions are seemingly regulated by proteolysis: S100A9 is readily degraded, while S100A8/S100A9 is resistant. We take an evolutionary biochemical approach to show that S100A9 evolved both functions and lost proteolytic resistance from a weakly proinflammatory, proteolytically resistant amniote ancestor. We identify a historical substitution that has pleiotropic effects on S100A9 proinflammatory activity and proteolytic resistance but has little effect on S100A8/S100A9 antimicrobial activity. We thus propose that mammals evolved S100A8/S100A9 antimicrobial and S100A9 proinflammatory activities concomitantly with a proteolytic “timer” to selectively regulate S100A9. This highlights how the same mutation can have pleiotropic effects on one functional state of a protein but not another, thus facilitating the evolution of multifunctionality.

## INTRODUCTION

The innate immune system uses a small number of multifunctional proteins to respond to diverse immune challenges.^1–4^ Multifunctional immune proteins are critical for pathogen defense,^2–4^ shaping host-associated microbial communities,^5^ and well-regulated tissue growth.^6–8^ They also drive pathological inflammation in disease, including autoimmune disorders, cancer, and cardiovascular disease.^9–13^ These multifunctional proteins raise both mechanistic and evolutionary questions. How can one protein sequence satisfy the multiple constraints imposed by having multiple functions? How can multiple functions evolve in one protein when, as a result of multifunctionality, each mutation likely has pleiotropic effects?^1, 14–16^

One such multifunctional protein is S100A9 (A9), a small, soluble protein found at high concentrations in the extracellular space during an inflammatory response.^17^ It has at least two key immune functions. As a homodimer, A9 potently activates inflammation via Toll-like receptor 4 (TLR4).^18–30^ As a heterocomplex with S100A8 (A8/A9, also known as calprotectin – figure 1a) it is antimicrobial.^31–44^ A9 exacerbates endotoxin-induced shock in mice.^45^ Both A9 and A8/A9 are primary biomarkers for many human inflammatory diseases.^46–50^ Further, dysregulation of A9 is associated with various cancers, pulmonary disorders, and Alzheimer’s disease.^19, 46–52^ Understanding the mechanisms by which A9 performs its innate immune functions is critical for developing treatments for A9-mediated diseases.

**Figure 1.**
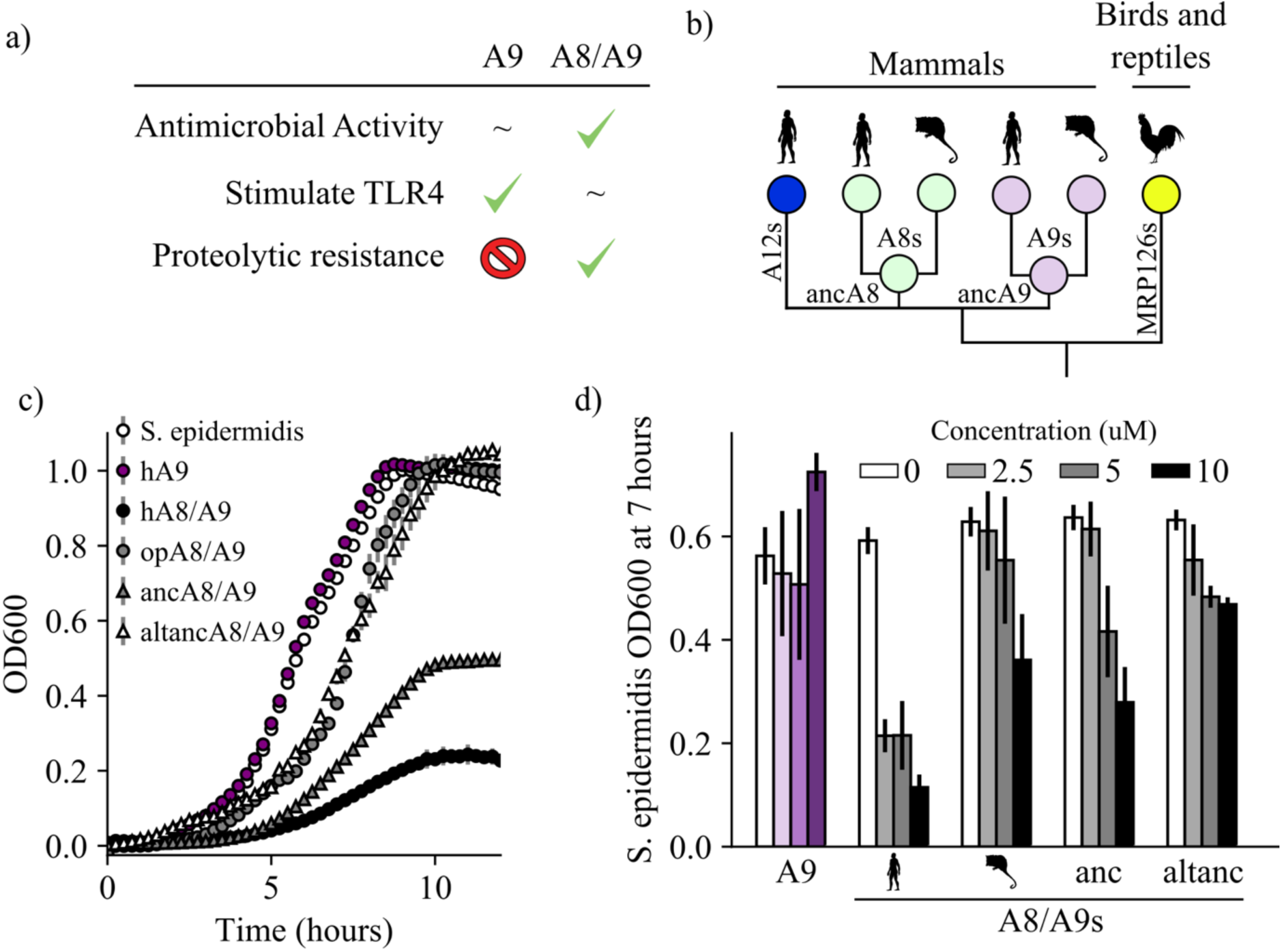
A9s evolved to form the antimicrobial A8/A9 complex early in mammals. (a) Table of A9 and A8/A9 properties. “∼” represents weak or ambiguously characterized function, check marks and red “X” represent confirmed property (check) or lack thereof (“x”). (b) Schematic of previously published S100 protein tree. Colored nodes represent single protein sequences. Species cartoons shown are human, opossum, and chicken. Ancestrally reconstructed protein nodes are labeled. Branch lengths not to scale. (c) Representative growth curves for *Staphylococcus epidermidis* in the presence or absence of 10 μM S100 proteins. Each point represents optical density at 600 nm. *S. epidermidis* growth alone and in the presence of modern proteins are shown as circles, growth in the presence of ancestrally reconstructed proteins shown as triangles. Error bars are standard deviation of three technical replicates. (d) Percent of untreated *S. epidermidis* growth at 12 hours with S100 protein treatments. Data are average of three biological replicates. Error bars are standard error of the mean. Species cartoon labels are the same as in (b).

The mechanism of A8/A9 antimicrobial activity is well established: it sequesters transition metals through a unique hexahistidine metal binding site at the A8/A9 heterodimer interface, thereby limiting the concentrations of essential microbial nutrients in the extracellular space.^31–44^ In contrast, the proinflammatory mechanism of A9 is not well understood. A9 acts as a Damage-Associated Molecular Pattern (DAMP), activating NF-κB and other cytokines through Toll-like receptor 4 (TLR4).^18–30^ The interaction interface(s), affinity, and stoichiometry for the A9/TLR4 interaction are not known. A small region of A9 has been suggested to form part of the A9/TLR4 binding surface,^53^ but no mutant of A9 has been identified that substantially compromises its activation of TLR4.

An additional layer of A9 immune function is that A9 and A8/A9 are thought to be regulated in the extracellular milieu by proteases. A9 is very susceptible to proteolytic degradation, while A8/A9 is highly resistant (figure 1a).^54, 55^ Proteolysis may serve to purge proinflammatory A9 from the extracellular space and thus selectively enrich for antimicrobial A8/A9. There may even be a direct, functional link between inflammation and proteolysis, as proteolytic fragments of A9 have been suggested to activate TLR4.^53^ Testing these ideas, however, has been challenging. It has been proposed that the origin of proteolytic resistance of A8/A9 is through the formation of a tetrameric dimer of heterodimers— (A8/A9)_2_, which forms in the presence of calcium.^56^ There is no obvious way, however, to selectively alter the proteolytic resistance of A9, making it difficult to test for correlation between A9 proteolytic resistance and A9 proinflammatory activity.

We took an evolutionary biochemical approach to mechanistically dissect the evolution of A9 innate immune functions. Using phylogenetics, ancestral sequence reconstruction (ASR), and biochemical studies, we show that A9s evolved to form proteolytically resistant, antimicrobial A8/A9 complexes in early mammals. We find that A9s gained proinflammatory activity and lost proteolytic resistance in the ancestor of therian mammals from a weakly proinflammatory, proteolytically resistant amniote ancestor. We identify a pleiotropic substitution that is necessary for A9 activation of TLR4, sufficient to increase TLR4 activation by the A9 amniote ancestor and played a role in loss of A9 proteolytic resistance. Mutating this site has minimal effect on A8/A9 antimicrobial activity or proteolytic resistance. Lastly, we show that proteolysis is not required for A9 activation of TLR4. Taken together, this work reveals that mammals concomitantly evolved A8/A9 antimicrobial activity, A9 proinflammatory activity, and a way to selectively regulate A9 inflammation via loss of A9 proteolytic resistance. These findings provide unprecedented mechanistic and evolutionary insight into A9 function and show how a single mutation can have pleiotropic effects in one functional state of a protein while not impacting another, thus facilitating the evolution of multifunctionality.

## RESULTS

We first set out to establish when A9 evolved three innate immune properties: antimicrobial activity via formation of the A8/A9 complex, proinflammatory activation of TLR4 by A9 alone, and the differential proteolytic susceptibility of A9 and A8/A9.

### A9s evolved to form antimicrobial A8/A9 complexes early in mammals

We sought to determine when A9 evolved to form the antimicrobial A8/A9 complex. We hypothesized that A8/A9 antimicrobial activity evolved in the ancestor of therian mammals (the shared ancestor of marsupials and placental mammals) for several reasons. First, the broad-spectrum antimicrobial activity of human and mouse A8/A9 is well established.^32–44^ Second, A9 and A8 genes are only found together in therian mammals (figure 1b);^57^ therefore the A8/A9 complex could not have arisen earlier than in the ancestor of therian mammals. Lastly, the residues composing the antimicrobial hexahistidine metal binding site are fully conserved across therian mammals (figure S1).

To determine whether the antimicrobial A8/A9 complex arose in the ancestor of therian mammals, we compared human A8/A9 to two previously uncharacterized A8/A9 complexes. We first tested the antimicrobial activity of A8/A9 from opossum, which is one of the earliest-diverging mammals relative to humans that possesses both of the S100A8 and S100A9 genes. Following previous work,^41^ we produced a cysteine-free variant of the complex to avoid the use of reducing agents in the antimicrobial assay. We confirmed that cysteine-free opossum A8/A9 formed a heterotetramer (46.8 ± 0.7 kDa) in the presence of calcium – like the human and mouse proteins^58^ – using size exclusion chromatography coupled with multi-angle laser light scattering (SEC MALS, figure S2).

**Figure 2.**
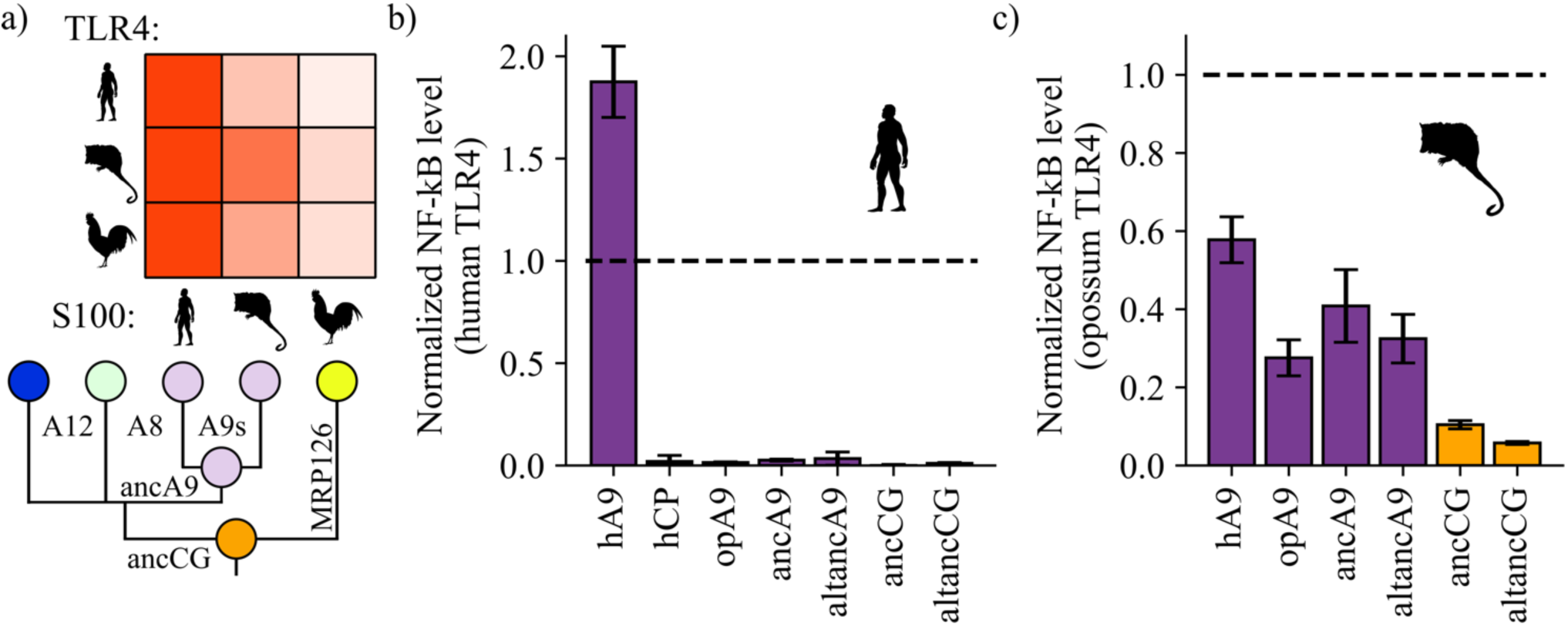
A9s gained proinflammatory activity from a weakly proinflammatory ancestor. (a) Schematic of previously measured proinflammatory activity of S100s against various TLR4s. Species labels on x and y-axes of heatmap are the same as figure 1. Heatmap coloring is scaled to match 2 μM S100 activity levels measured in supplementary figure S2 of Loes et al. 2018.^57^ (b) and (c) NF-κB production of human and opossum TLR4 in response to treatment with modern and ancestral S100 proteins. Bars represent average of >3 biological replicates, error bars are standard error of the mean. All values are background-subtracted and normalized to LPS positive control (see methods).

We measured cysteine-free opossum A8/A9 antimicrobial activity against *Staphylococcus epidermidis* using a previously established *in vitro* antimicrobial assay that monitors bacterial growth in the absence or presence of S100 proteins (figure 1c, S3).^41^ To compare the activity of different proteins, we quantified inhibition at seven hours (figure 1d). We observed a dose-dependent decrease in *S. epidermidis* growth in the presence of low micromolar concentrations of cysteine-free opossum A8/A9 (figure 1c-d, S3). The antimicrobial activity of opossum A8/A9 was weaker than that of human A8/A9: opossum A8/A9 delayed bacterial growth, while human A8/A9 both delayed growth and decreased bacterial carrying capacity (figure 1c). It was previously found that cysteine-free human A8/A9 was potently antimicrobial,^41^ while cysteine-free mouse A8/A9 exhibits weaker antimicrobial activity than wildtype mouse A8/A9.^32^ To determine whether the weaker activity of opossum A8/A9 was due to the removal of cysteines, we also measured the activity of wildtype opossum A8/A9 against *S. epidermidis*. We found that wildtype opossum A8/A9 had higher activity than cysteine-free opossum A8/A9 over the concentration range tested (figure S3). This suggests that the cysteines present in mouse and opossum A8/A9 play a role in their antimicrobial activity, unlike in human A8/A9. The antimicrobial activity of the opossum A8/A9 complex thus appears to be more similar to that of mouse A8/A9 than human A8/A9.

**Figure 3.**
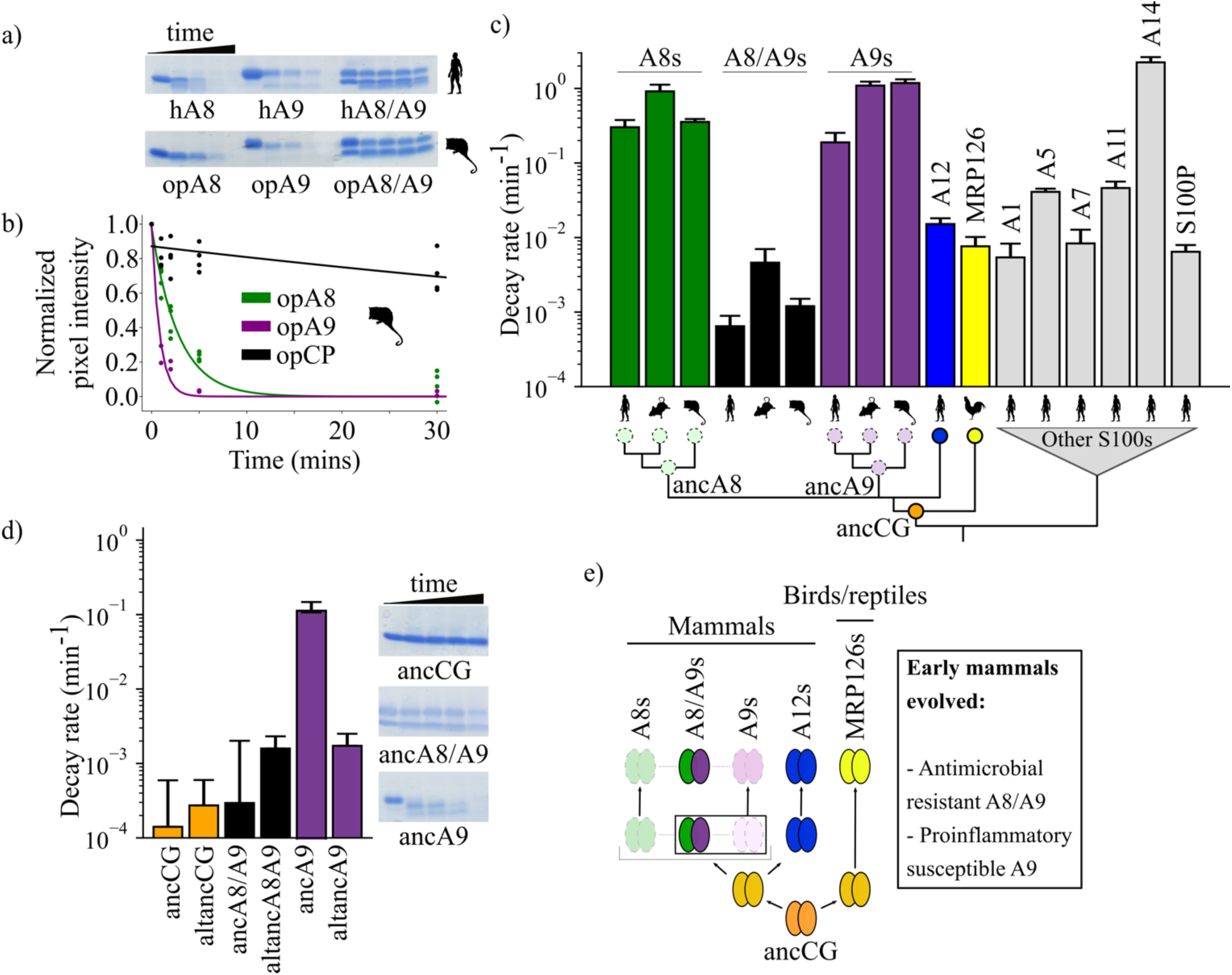
A9s lost proteolytic resistance from a proteolytically resistant amniote ancestor. (a) *In vitro* proteolytic resistance assay showing SDS-PAGE gel of S100 protein degradation via proteinase K over time. Gels were quantified using densitometry and normalized to the undigested protein band intensity. (b) A single exponential decay model was globally fit to the data to quantify decay rates. Points are biological replicates, lines are model fit to data. (c) S100 protein proteolysis rates mapped onto schematized S100 phylogeny. X-axis cartoon labels same as in figure 1. Circles indicate proteolytic susceptibility (faded/dashed) and resistance (solid), with predicted resistance shown for ancA8, ancA9, and ancCG nodes. (d) Decay rates for ancestrally reconstructed proteins, with gels shown on the right. For panels (c) and (d), error bars are the square root of the diagonalized covariance matrix from the fit and the y-axis is in log scale. (e) Summary model for proposed evolution of A9 and A8/A9 innate immune properties. Box around A8/A9 and A9 indicate location in tree (ancestor of therian mammals) where immune functions evolved.

The shared antimicrobial activity of human, mouse, and opossum A8/A9 strongly suggests that the antimicrobial A8/A9 complex evolved in the ancestor of therian mammals. To test this further, we measured the antimicrobial activity of ancestrally reconstructed therian mammalian A8/A9 (ancA8/A9 – figure 1b). We used our previously published phylogenetic tree^57^ consisting of 172 S100 sequences to reconstruct therian mammalian ancestral A8 and A9 (ancA9 and ancA8 – figure S4), which were used to form the ancA8/A9 complex. These proteins had average posterior probabilities of 0.83 and 0.88, respectively (table S4). We confirmed that each protein was folded and had secondary structure content similar to that of human A8/A9 using far-UV circular dichroism (CD) spectroscopy (figure S5).

**Figure 4.**
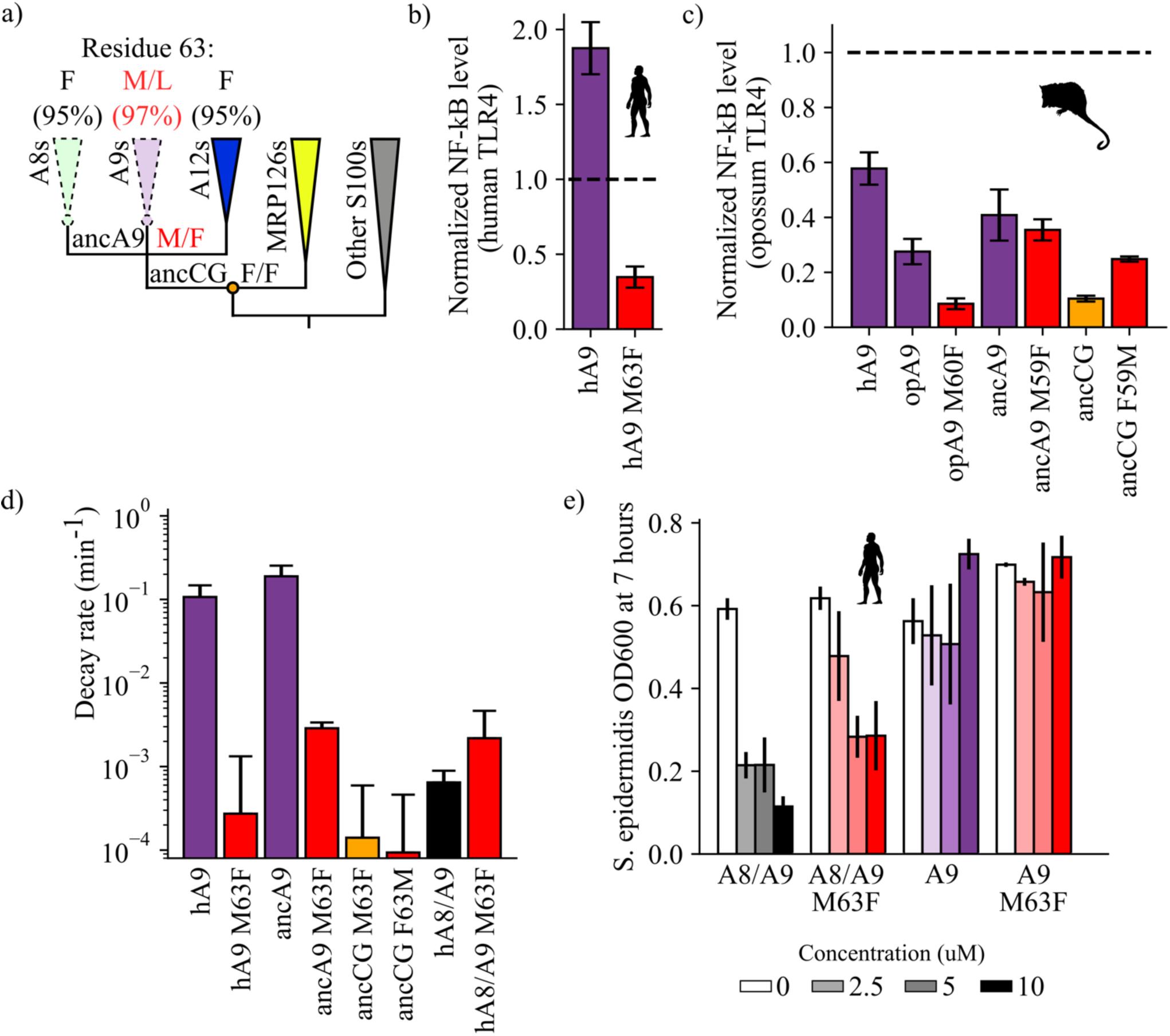
A single historical substitution affects A9 proinflammatory activity and proteolytic resistance without affecting A8/A9 proteolytic resistance or antimicrobial activity. (a) Schematic S100 phylogenetic tree with the amino acid state of position 63 shown at key nodes. Wedges represent clades, colored as in figure 1. Lines indicate proteolytic susceptibility (faded/dashed) and resistance (solid). Circles indicate characterized ancestors. Amino acid labels represent maximum likelihood state/alternate amino acid state for position 63 at ancestral nodes, while labels at clade tips represent percent conservation across modern S100 protein sequences. (b-c) NF-κB production of S100 point mutants at position 63 against human (b) and opossum (c) TLR4. (d) Proteolysis rates for S100 point mutants at position 63. Error bars and y-axis are the same as in figure 1. (d) Antimicrobial activity of hA9 and hA8/A9 with and without M63F mutation against *S. epidermidis.* Axes and error bars same as in figure 1d.

**Figure 5.**
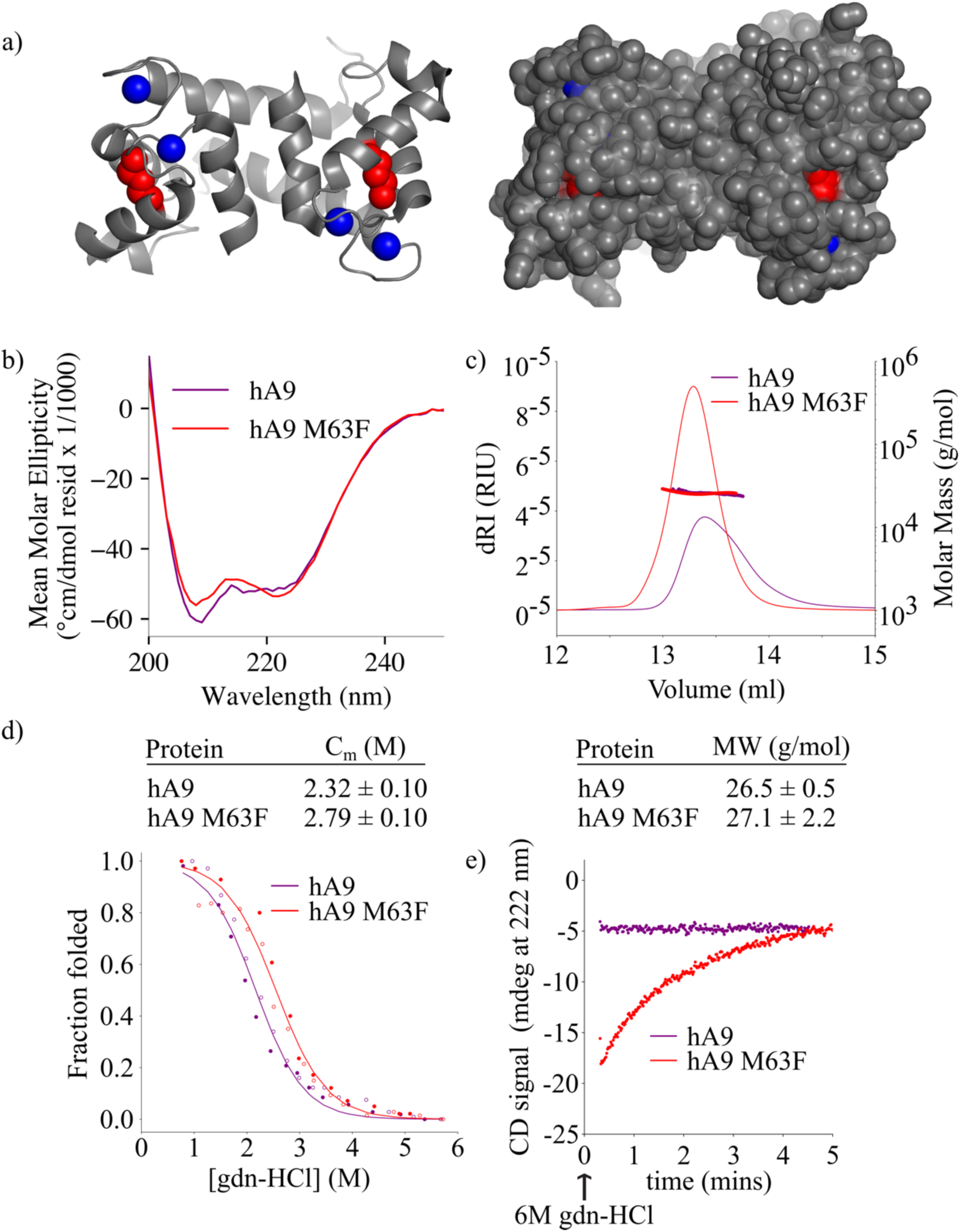
M63F increases human A9 apparent stability by decreasing its unfolding rate. (a) Crystal structure of hA9 (PDB entry 1irj).^68^ Cartoon depiction left, surface view right. Calcium ions are blue spheres. M63 is highlighted in red – two total for homodimeric A9. (b) Far-UV circular dichroism (CD) spectroscopy scans of hA9 and hA9 M63F. Data represent average of 3 scans. (c) SEC MALS analysis of hA9 and hA9 M63F oligomeric state. Solid lines are refractive index (left y-axis), points and molecular weights in table below represent molar mass calculated from light scattering detectors using ASTRA software (right y-axis - see methods). (d) Equilibrium chemical denaturation of 0.5 uM hA9 and hA9 M63F monitored by CD at 222 nm. Filled circles are protein unfolding, empty circles are refolding. A two-state denaturation model was fit to the data to extract the Cm – errors in table below represent standard error of the mean of three biological replicates (p = 0.02; paired t-test) (figure S14). (e) Kinetics of hA9 and hA9 M63F unfolding via chemical denaturation. Graph depicts one representative unfolding experiment (figure S15).

We then measured the antimicrobial activity of ancA8/A9 against *S. epidermidis.* We observed a potent reduction in *S. epidermidis* growth comparable to that of human A8/A9 (figure 1c-d, S3). To test for the robustness of this finding to phylogenetic uncertainty, we also tested the antimicrobial activity of an AltAll^59^ reconstruction of ancA8/A9 against *S. epidermidis* (altancA8/A9, figure 1c-d, S3). In this reconstruction, we swapped all sites with ambiguous reconstructions for their second-most likely state (see methods). AncA8/A9 and altancA8/A9 differ by 24 amino acids. AltancA8/A9 exhibited antimicrobial activity against *S. epidermidis* similar to opossum A8/A9: it delayed growth but did not ultimately limit bacterial carrying capacity. While the hexahistidine site residues are conserved in ancA8/A9 and altancA8/A9 (figure S1), it appears that a subset of the ambiguously reconstructed 24 residues are important for A8/A9 antimicrobial activity, perhaps affecting the orientation and/or affinity of the hexahistidine metal binding site.

Taken together, the antimicrobial activity of modern mammalian A8/A9 complexes (human, mouse, and opossum) and the antimicrobial activity of the reconstructed ancA8/A9 complex suggest that A9s evolved to form the antimicrobial A8/A9 complex in the ancestor of mammals.

### A9s evolved potent proinflammatory activity from a weakly proinflammatory amniote ancestor

We next sought to determine when A9s evolved potent proinflammatory activity via activation of TLR4. Our previous work revealed that human A9 potently activates not only human TLR4 in functional assays, but also opossum and chicken TLR4 (figure 2a).^57^ In contrast, chicken MRP126, the sauropsid ortholog of A9s, was found to be a weak activator of all TLR4s, including chicken TLR4. Both human and opossum A9 activate chicken TLR4 better than chicken MRP126 does. Two possibilities are consistent with these observations. Either mammalian A9s evolved enhanced proinflammatory activity from a less active amniote ancestral state, or A9s maintained a potent ancestral activity that was lost by chicken MRP126.

To differentiate between these two possibilities, we determined the ancestral proinflammatory activity of these proteins. We used ASR to reconstruct the shared amniote ancestor of A9s, A8s, A12s, and MRP126s. This group of proteins is known collectively as “calgranulins”, so we will refer to this ancestral protein as ancCG (ancestor of calgranulins). We also constructed an alternate, “alt All” version of this ancestor (altancCG, S4). The average posterior probability of ancCG was 0.86 (table S4). We also expressed and purified ancA9 and altancA9 – the A9 subunits from the ancestral A8/A9 complexes described above. We confirmed that each protein was folded and had secondary structure content similar to that of modern S100s using far-UV CD spectroscopy (figure S5).

We then tested modern and ancestral S100s for activity against human TLR4. Following previous work,^45, 57, 60, 61^ we transiently transfected HEK293T cells with plasmids encoding TLR4 and its species-matched cofactors MD-2 and CD14, added purified S100 proteins to the growth media, and then measured output of luciferase under control of an NF-κB promoter. Consistent with previous results,^57^ we found that human A9 potently activated human TLR4, resulting in high levels of NF-κB production (figure 2b, S6-7). Human A8/A9 and opossum A9 exhibited much weaker activity against human TLR4. Lastly, we tested ancA9, altancA9, ancCG, and altancCG for activity against human TLR4 and observed weak or no activation for each ancestral protein. This result is unsurprising, as we previously found that human TLR4 is more specific than other amniote TLR4s: human TLR4 is activated much more potently by human A9 than by any other S100 protein (figure 2a-b).^57^ In contrast, TLR4s from other species (mouse, opossum, and chicken) appear to be more promiscuous and can be activated similarly by S100s from various species (figure 2a).^57^ This is consistent with lineage-specific coevolution between human TLR4 and human A9 – a confounding variable that makes assessment of ancestral S100 protein proinflammatory activity difficult using human TLR4.

**Figure 6.**
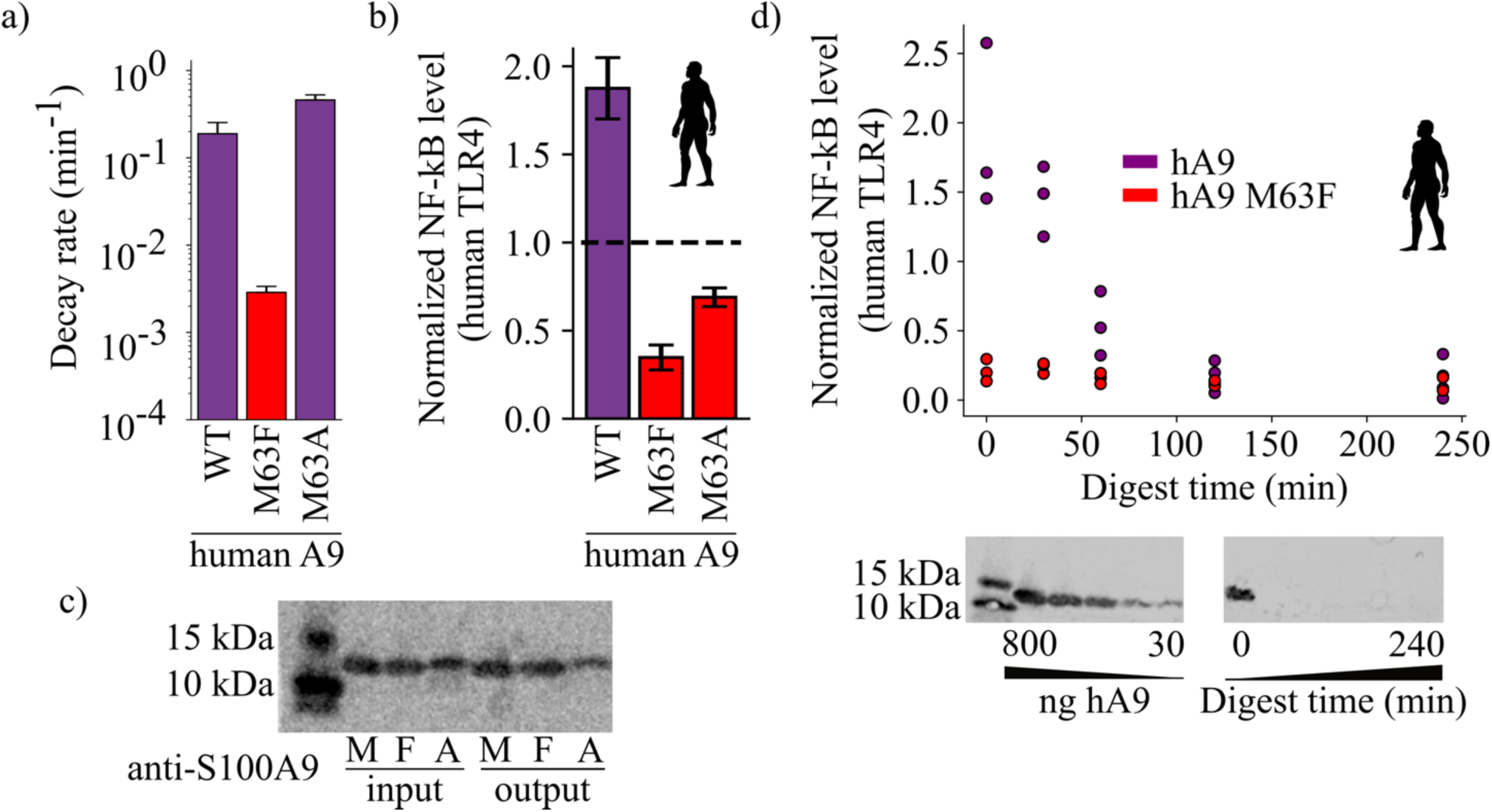
Proteolysis is not required for A9 activation of TLR4. (a) Proteolytic decay rates for human point mutants at position 63. Error bars and axes are the same as in figure 3. (b) NF-κB production of human TLR4 in response to treatment with hA9, hA9 M63F, and hA9 M63A. Error bars the same as in figure 2. (c) Western blot of hA9 and position 63 point mutants before and after proinflammatory activity assay. Left bands represent 10 and 15 kDa ladder. (d) NF-κB production of human TLR4 in response to hA9 and hA9 M63F pre-proteolyzed with proteinase K for increasing amounts of time. Points are biological replicates and are the average of three technical replicates. Western blots below depict the amount of full-length A9 remaining over time. Left blot shows antibody sensitivity to A9, right shows digestion time course samples. Ladder and antibody same as in (c).

We predicted that opossum TLR4 would be a better protein to probe ancestral S100 proinflammatory function because opossum TLR4 is broadly activated by A9s across mammals and gives little indication of lineage-specific coevolution.^57^ We therefore tested the proinflammatory activity of ancA9, ancCG, and their corresponding alternate reconstructions against opossum TLR4. Corroborating previous results, human A9 strongly activated opossum TLR4, while opossum A9 activity was approximately half that of human A9 (figure 2c, S6-8). AncA9 and altancA9 activated opossum TLR4 to the same extent as opossum A9. AncCG and altanCG were the weakest activators of opossum TLR4, with activity approximately 25% or less than that of human A9.

These findings suggest that A9s evolved enhanced proinflammatory activity early in mammals from a weakly proinflammatory amniote ancestor, while A8/A9s and chicken MRP126 maintained weak, ancestral proinflammatory activity.

### A9s evolved proteolytic susceptibility from a proteolytically resistant amniote ancestor

We next sought to determine when the differential proteolytic susceptibility of A9 and A8/A9 evolved. We used a simple *in vitro* assay to monitor S100 protein degradation over time in the presence of proteinase K, a potent non-specific serine protease (figure 3a). Proteinase K was chosen both because of its low specificity and to mimic other serine proteases that A9 and A8/A9 encounter when released from neutrophils during an inflammatory response.^62–66^ Proteolytic decay rates were estimated by fitting a single exponential decay function to the data (figure 3b, S9-12).

Human A8/A9 has been described as extremely resistant to proteases.^54^ however, it has not been compared to S100 proteins besides human S100A8 (A8) and A9. To establish a baseline expectation for S100 protein proteolytic resistance, we characterized the proteolytic resistance of a broad set of human S100s against proteinase K. As previously shown,^54^ human A9 and A8 alone were rapidly proteolytically degraded while the human A8/A9 complex exhibited strong resistance (figure 3c, S9-10). Under our conditions, the degradation rates for human A8 and A9 were approximately three orders of magnitude faster than that of the human A8/A9 heterocomplex. We then characterized closely related protein human S100A12 (A12), the chicken ortholog MRP126, and six distantly related human S100s.^67^ Human A12, chicken MRP126, and five out of six more distantly related human S100s exhibited intermediate to strong proteolytic resistance, each degrading 1-2 orders of magnitude slower than human A8 or A9 but, on average, one order of magnitude faster than human A8/A9 (figure 3c, S9). Notably, human A12 and chicken MRP126 formed predominantly homodimers by SEC MALS under these conditions (figure S2), indicating that higher-order oligomerization (> 2 subunits) isn’t required for S100 proteolytic resistance. Lastly, human A14 degraded faster than A9 or A8. This protein is evolutionarily distant^67^ and therefore likely reflects independent evolution of this property. Taken together, these data show that the A8/A9 complex, A9, and A8 indeed fall at the extremes of human S100 proteolytic resistance; human A9 and A8 are among the fastest-degrading S100s tested, while human A8/A9 is one of the slowest.

To test whether A9 and A8 proteolytic susceptibility and A8/A9 resistance are conserved across mammals, we characterized mouse and opossum A9, A8, and A8/A9 for proteolytic resistance. Mouse A9 and A8 were found to be highly proteolytically susceptible and mouse A8/A9 strongly proteolytically resistant, matching the pattern observed for their human counterparts (figure 3c and S10). Opossum A9 and A8 were also highly proteolytically susceptible, while opossum A8/A9 was resistant (figure 3c, S10). This indicates that the susceptibility of A9s and A8s and the resistance of A8/A9 complexes is conserved across mammals.

When mapped onto the S100 phylogeny, the most parsimonious explanation for these data is that the shared amniote ancestor—ancCG—was proteolytically resistant (figure 3c). In this scenario, A12s, MRP126s, and A8/A9s conserved ancestral resistance, while A9s and A8s independently lost resistance early in mammals. Alternatively, ancCG could have been proteolytically susceptible. This would mean that A9s and A8s maintained an ancestral susceptibility, while MRP126s, A12s, and A8/A9s each evolved novel proteolytic resistance.

To distinguish between these possibilities, we characterized ancestrally reconstructed S100s for proteolytic resistance. AncCG and altancCG exhibited extremely high proteolytic resistance (figure 3d, S11), with degradation rates 3-4 orders of magnitude slower than modern A8s or A9s and approximately one order of magnitude slower than modern A8/A9s. AncA8/A9 and altancA8/A9 also demonstrated high proteolytic resistance, with degradation rates approximately 2-3 orders of magnitude slower than A8s and A9s and comparable to modern A8/A9 complexes (figure 3d, 2f). Together, these data paint a consistent picture: the amniote ancestor of A9s, ancCG, was strongly resistant to proteolytic degradation. Modern A9s and A8s lost proteolytic resistance from an ancestrally resistant state, while modern A12s, A8/A9 complexes, and MRP126s maintained the ancestral proteolytic resistance (figure 3e).

Finally, we sought to better resolve when A9s acquired proteolytic susceptibility. We hypothesized that this occurred in the ancestor of mammalian A9s before the divergence of therian mammals and marsupials. To test this hypothesis, we measured the proteolytic susceptibility of therian mammalian ancA9 and found that it degraded rapidly. However, its alternative reconstruction (altancA9), was slow to degrade, with a rate two orders of magnitude slower than ancA9 and comparable to other highly resistant S100s. Because the descendants of ancA9 all exhibit proteolytic susceptibility, the simplest explanation is that altancA9 is a low-quality reconstruction that does not capture the properties of the historical protein. Alternatively, proteolytic susceptibility could have been independently acquired along marsupial and placental mammal lineages.

### A single substitution had pleiotropic effects on TLR4 activity and proteolytic resistance

We found above that A9 evolved to form the antimicrobial A8/A9 complex, gained potent proinflammatory activity, and lost proteolytic resistance over the narrow evolutionary interval after the divergence of mammals and sauropsids but before the divergence of placental mammals and marsupials. We next sought to determine how A9 evolved its antimicrobial and proinflammatory activities and lost proteolytic resistance.

The mechanism by which A9 evolved to form the antimicrobial A8/A9 complex appears straightforward. After the duplication that gave rise to A8 and A9, mutations accumulated in both molecules that created the antimicrobial hexahistidine metal binding site in the A8/A9 complex (figure 3d). Mammalian A9s also have a conserved C-terminal extension that contributes two histidines to the hexahistidine metal binding site. The quantitative difference between ancA8/A9 and altancA8/A9 antimicrobial activity suggests that other amino acid changes tuned the antimicrobial activity of the molecule, but the core functionality is determined by whether the six histidine residues were present.

The mechanisms by which A9s gained proinflammatory activity and lost proteolytic resistance are less obvious, particularly because the mechanism by which A9 activates TLR4 is not well understood. We reasoned that we could identify functionally important amino acid substitutions by focusing on the evolutionary interval over which these properties evolved. We therefore compared the sequences of ancCG (weakly proinflammatory and resistant to proteolytic degradation) and ancA9 (potently proinflammatory and susceptible to proteolytic degradation). We further narrowed down sequence changes of interest by looking for residues conserved in modern A9s (figure S13). Finally, we focused on amino acid changes in helix III of A9, as this region is thought to be important for A9 activation of TLR4 based on *in vitro* binding studies and *in silico* docking studies.^53^ Only one historical amino acid substitution met all three criteria: position 63 (human A9 numbering). This residue is a phenylalanine in both ancCG and altancCG, is conserved as a phenylalanine in 95% of modern A8s and A12s and has been substituted for a methionine or leucine (M/L) in 97% of A9s (figure 4a).

We hypothesized that reverting this site to its amniote ancestral state—M63F—might affect A9 proinflammatory activity. We mutated this position to a phenylalanine in human A9 and opossum A9 and tested each protein for TLR4 activation. Strikingly, we found that introducing M63F into human A9 severely compromised its ability to activate human TLR4 (figure 4b, S7). This was also true for opossum A9: introduction of M63F strongly decreased opossum A9 activation of opossum TLR4 (figure 4c, S8). We next introduced the forward substitution, F63M, into ancCG and tested its proinflammatory activity against opossum TLR4. We observed a modest increase in ancCG activity with the F63M substitution, with activity comparable to that of opossum A9 (figure 4c, S8).

For most proteins we studied, the amino acid at position 63 did indeed play an important role in determining the pro-inflammatory activity of A9. The effects of toggling position 63 between Met and Phe were not, however, universal. We introduced M63F into ancA9 and observed no change in proinflammatory activity (figure 4c, S8). Further, altancA9 has a Phe at position 63 but activates TLR4 in the assay (figure 2c, S8). Thus, while position 63 is an important contributor to activity in modern A9s, other substitutions were also important for the transition from a weakly pro-inflammatory ancestor to the modern set of potently pro-inflammatory A9s.

Because A9s lost proteolytic resistance and gained proinflammatory activity over the same evolutionary time interval, we reasoned that the F63M substitution might have also played a role in A9 loss of proteolytic resistance. To test this, we characterized the proteolytic resistance of human A9 M63F and ancA9 M63F. Strikingly, reversion of this single mutation rendered both ancA9 and human A9 strongly resistant to proteolytic degradation, decreasing their respective degradation rates by 1-2 orders of magnitude and approaching the degradation rates of ancCG and various A8/A9 complexes (figure 4d, S11). We also tested the effect of the forward mutation – F63M – on ancCG proteolytic resistance. We observed no change in resistance for ancCG F63M, indicating that additional substitutions were required to render ancA9 proteolytically susceptible. Together these data show that a single historical reversion is sufficient to render A9s proteolytically resistant, indicating that this position played a role in the loss of A9 proteolytic resistance early in therian mammals.

### The pleiotropic substitution does not affect antimicrobial activity

We then investigated whether introducing M63F affects the properties of the antimicrobial A8/A9 complex. As A9 is a critical subunit of A8/A9, we reasoned that perhaps position 63 was altered in A9s due to some incompatibility between Phe63 and A8/A9 formation or function. To this end, we introduced M63F into human A8/A9 and tested it for oligomeric state, proteolytic resistance, and antimicrobial activity against *S. epidermidis.* Human A8/A9 M63F predominantly formed a heterotetramer in the presence of calcium by SECMALS with a molecular weight similar to that of wildtype human A8/A9 (48.7 ± 4.2 kDa – figure S2). We found that human A8/A9 M63F was also strongly resistant to proteolytic degradation, similar to human A8/A9 (figure 4d). Lastly, M63F had minimal impact on human A8/A9 antimicrobial activity against *S. epidermidis*, retaining potent antimicrobial activity (figure 4e). In contrast, neither human A9 nor human A9 M63F were antimicrobial against *S. epidermidis* (figure 4e). These findings suggest that this single amino acid position had important effects on the evolution of A9 activation of TLR4 and loss of proteolytic resistance without significantly impacting A8/A9 oligomeric state, proteolytic resistance, or antimicrobial activity.

### M63F increases protein thermodynamic stability and decreases unfolding rate of human A9

We next asked what effect M63F has on the biophysical properties of human A9. Residue 63 sits in the middle of helix III of A9, pointing inward toward helix II, and is neither a core residue nor fully surface-exposed (figure 5a).^68^ Based on the published structure of human A9,^68^ a Phe at position 63 could be plausibly tolerated without a steric clash. Using circular dichroism (CD) spectroscopy, we found that the bulk secondary structure content of human A9 M63F was similar to that of hA9 (figure 5b). We measured the oligomeric state of human A9 M63F by SEC MALS and found that it predominantly forms a homodimer in solution similarly to human A9, with no detectable monomers or larger oligomers (figure 5c). These data together indicate that M63F doesn’t significantly alter human A9’s secondary structure or oligomeric state.

We then examined whether M63F alters the stability of human A9. We measured unfolding and refolding curves for human A9 and human A9 M63F using CD spectroscopy and chemical denaturation via guanidine hydrochloride (gdn-HCl). We found that M63F appears to stabilize human A9, shifting the C_m_ by approximately 0.5 M (figure 5d, S14). We also measured the unfolding kinetics of human A9 and human A9 M63F in the presence of calcium by spiking protein directly into 6M gdn-HCl and monitoring its unfolding rate by CD spectroscopy. Strikingly, human A9 M63F takes several minutes to unfold under these conditions, while human A9 unfolds immediately within the dead time of the experiment (figure 5e, S15). We note that the folding pathway for A9 is complex and almost certainly not two-state—calcium binding, monomer folding, and dimerization all contribute—and thus we cannot reliably determine how M63F affects the stability of each of these potential folding intermediates. The large increase in the apparent unfolding rate reveals, however, that the mutation stabilizes some aspect of the folded structure rather than simply increasing the rate of folding, oligomerization and/or calcium binding.

### Proteolysis is not required for A9 activation of TLR4

The work above identified a mutation that, when introduced into human A9, increases the stability of the protein structure while also potently compromising the ability to activate TLR4. The mutation is not at a surface position and is therefore not likely a direct participant in the A9/TLR4 protein/protein interface. Further, the same mutation dramatically decreases the proteolytic susceptibility of the protein. One simple way to explain these observations would be if the proteolytic susceptibility itself was the feature that evolved to allow activation of TLR4. This would be consistent with a previous observation that proteolytic products of A9 activate TLR4.^53^

To test whether proteolysis itself was sufficient for activity, we engineered an alternate variant of A9 that was proteolytically susceptible. We introduced the M63A mutation into human A9, anticipating that the short alanine sidechain would not have the stabilizing effect of M63F. As expected, human A9 M63A was highly susceptible to proteolytic degradation, similar to wildtype human A9 (figure 6a, S12).

We reasoned that if proteolysis is the primary determinant of A9 activation of TLR4, then proteolytically susceptible human A9 M63A should potently activate TLR4. Human A9 M63A, however, exhibited diminished proinflammatory activity, similar to human A9 M63F (figure 6b, S7). This indicates that the methionine at position 63 is important for A9 activation of TLR4. Further, we quantified the amount of human A9, human A9 M63F, and human A9 M63A before and after measuring TLR4 activity and observed no decrease in the amount of full-length protein remaining for wildtype human A9 or either mutant by western blot (figure 6c). This indicates that A9 is not digested by extracellular proteases over the course of the *ex vivo* assay and that proteolysis is likely not necessary for A9 activation of TLR4.

Although proteolysis does not appear to be a requirement for TLR4 activation, this does not rule out that proteolysis could increase A9 proinflammatory activity by releasing proinflammatory fragments of A9. To test for this possibility, we treated human A9 with agarose-immobilized proteinase K for increasing amounts of time, removed the protease, and then measured the proinflammatory activity of A9 degradation products (figure 6d). If proteolytic products of A9 are the most proinflammatory form of the protein, we might expect to observe a spike in TLR4 activation upon A9 digestion. Instead, we observed a steady decrease in human A9 activity with increasing digestion time. This suggests that full-length human A9 is the most potent activator of TLR4.

We did observe moderate activity for proteolytic products of human A9, as previously shown.^53^ After 30 minutes of digestion, no detectable full-length A9 remains by western blot (< 30 ng, figure 6d), but NF-κB production is still quite high, revealing that smaller fragments of A9 are sufficient to provide some degree of activation of TLR4. This raised the possibility that part of M63F’s deleterious effect on proinflammatory activity could be to limit the release of active proteolytic fragments of A9. To test this, we also measured human A9 M63F activation of TLR4 after digestion for multiple hours (figure 6d). Unlike wildtype, however, fragments of human A9 M63F did not activate TLR4—even after being liberated by the protease. This strongly suggests that the historical mutation induced some change in the native structure or dynamics of the molecule to bring about increased activity, independent of its effect on proteolytic susceptibility.

## DISCUSSION

The work presented here provides insight into how the multifunctional protein A9 evolved critical innate immune functions. We find that mammalian A9s gained enhanced proinflammatory activity and lost proteolytic resistance from a resistant, weakly proinflammatory amniote ancestor. A single substitution played a key role in the evolution of these properties without significantly affecting the antimicrobial activity of the A8/A9 heterocomplex. This work contributes to our mechanistic understanding of how A9 activates TLR4 to drive inflammation and clarifies the role of proteolysis in A9 innate immune function.

### An evolutionary lens provides unique insight into the evolution of A9 innate immune functions

How did A9s evolve multifunctionality over a short evolutionary window? One might expect that the protein was under pleiotropic constraint to optimize each function.^1, 15, 16, 69–71^ However, we show that one amino acid position is pleiotropic for one functional state of A9 (the A9 homodimer) but not the other (the A8/A9 heterocomplex). Reverting M63F in human A9 has a pleiotropic effect on its proinflammatory activity and proteolytic resistance but has little effect on human A8/A9 antimicrobial activity or proteolytic resistance. This suggests that mammals benefitted from this pleiotropic substitution early in A9s by acquiring a proinflammatory molecule that can be regulated via proteolytic degradation without altering the critical antimicrobial function of the A8/A9 complex. This reveals that the same mutation can have pleiotropic effects on one functional state of a protein while leaving another untouched, thus providing an advantageous route for evolving multifunctionality.

This study highlights the utility of taking an evolutionary approach to study protein function.^72^ Previous work examining the proteolytic resistance of S100s focused on the extreme proteolytic resistance of A8/A9, which was first observed over ten years ago.^54^ Here we show that many S100s are, in fact, proteolytically resistant (figure 3c). By examining the proteolytic resistance of S100s through an evolutionary lens, we find that it is the loss of A9 resistance—not a gain of resistance in A8/A9—that was the important evolutionary change that occurred in these proteins. This approach facilitated identification of a single site (position 63)—out of more than 100—that strongly correlates with both A9 proteolytic susceptibility and A9 activation of TLR4. Isolating this position would have been very challenging without an evolutionary approach, as it does not stand out in a simple multiple sequence alignment or as part of the protein structure. We note that while reverting this site to its ancestral state is sufficient to render A9s resistant, further mutations were required between the amniote (ancCG) and mammalian A9 (ancA9) ancestors to render A9s susceptible, as introduction of F63M into ancCG did not decrease this protein’s proteolytic resistance (figure 4d).

Why did A9s lose proteolytic resistance? We suggest three possibilities. The first is that loss of proteolytic resistance in A9s was simply a byproduct of evolving proinflammatory activity. No A9 characterized in this study, with the exception of the alternate reconstruction of ancA9, is both proteolytically resistant and potently proinflammatory. This indicates that the molecular requirements for A9 proteolytic resistance may be incompatible with those required for A9 activation of TLR4, and A9s may have gained proinflammatory activity at the expense of proteolytic resistance. A second possibility is that A9 proteolytic susceptibility is being maintained to actively remove proinflammatory A9 from the cell and retain the antimicrobial A8/A9 complex. The last possibility for A9 loss of proteolytic resistance is adaptive constraint. There could be selection for some property of A9 or A8/A9 that we did not measure that is incompatible with A9 proteolytic resistance.

While we cannot explicitly distinguish between each of these possibilities, the end result is that A9s evolved potent proinflammatory activity while simultaneously evolving a proteolytic “timer” to regulate it, all without affecting A8/A9 function.

### Innate immune functions of A9 continued to evolve within the mammals

Our data suggest that the proinflammatory and antimicrobial activities of A9 and the A8/A9 complex have undergone further optimization in placental mammals since these functions evolved. While the histidines composing the high-affinity metal binding site of A8/A9 complexes are conserved, we observed differences in antimicrobial potency for different A8/A9 complexes. In particular, human A8/A9 is one of the most potently antimicrobial A8/A9 complexes characterized. This suggests that further optimization of the metal binding site has occurred in along the human lineage within mammals. We also observed differences in activation of TLR4 by different A9s—human A9 is a potent, promiscuous activator of TLR4s from multiple species, while earlier-diverging A9s and other S100s exhibit weaker proinflammatory activity.^57^ Future studies are necessary to understand how, mechanistically, later-diverging A9s and A8/A9 complexes have optimized these critical innate immune functions.

### Novel mechanistic insight into A9 activation of TLR4

Our findings suggest new directions for understanding how A9 potently activates TLR4. TLR4-driven inflammation has been the focus of intense study for over 20 years,^19, 23–25, 27, 60, 73–76^ and the structural basis of TLR4 activation by exogenous agonists, such as bacterial cell wall component lipopolysaccharide (LPS), is well understood.^77^ In contrast, very little is known about how A9 activates TLR4. We have shown here that proteolytic degradation appears dispensable for activation; however, smaller fragments of the protein are sufficient activate TLR4 (figure 6). Given the effect of mutating position 63 on A9 proinflammatory activity, we propose that the region surrounding it—helix III—is important for activity. This is independently supported by Vogl et al., who identified four pairs of double mutants within helix III (amino acids 64, 65, 73, and 77) that, when mutated to alanines in pairs, decrease A9 binding to TLR4 *in vitro*.^53^ Biophysical characterization of hA9 M63F (figure 5) indicates that it is more stable and unfolds more slowly, yet it maintains its bulk secondary structure and oligomeric state. The simplest explanation for these data is that M63F is affecting some functionally important dynamic process of the protein, possibly mediated by helix III, that is critical for A9 activation of TLR4. The proteolytic susceptibility of A9s also supports this hypothesis, as proteolysis is a dynamic process that often relies on substrate flexibility and local unfolding events to proceed.^78–82^ Damage-Associated Molecular Patterns (DAMPs) often interact with their targets via hydrophobic surfaces;^83–85^ one possibility is that A9 undergoes a local unfolding event that exposes a hydrophobic surface to interact with TLR4. This would mean that studies of the native structure of A9 might not be sufficient to gain mechanistic understanding of how it activates TLR4. Further work is required to understand the nature of the active functional state of A9.

### Pleiotropic mutations can facilitate the evolution of multifunctionality

Finally, our results suggest a positive role for pleiotropy in the evolution of protein function. Pleiotropy is often viewed as a constraint on evolution: as functional complexity is added to a polypeptide sequence, it becomes increasingly challenging to introduce substitutions—and new functions—without perturbing existing ones.^1, 15, 16, 69, 70, 86^ Here, however, we find a single mutation that had beneficial pleiotropic effects on two important properties of A9: proinflammatory activity and proteolytic susceptibility. If A9 evolved potent proinflammatory activity without gaining susceptibility, it could potentially overstimulate inflammation simply by lingering in the extracellular milieu. Since both properties evolved at once, however, mammals evolved a proinflammatory molecule with a built-in “timer”: they gained a new inflammatory signal while avoiding potentially deleterious effects. This shows how pleiotropy can positively contribute to the evolution of new functions.

This same mutation, in contrast, had little pleiotropic effect on another functional state of A9: the A8/A9 complex. The antimicrobial activity of the A8/A9 complex was insulated from any pleiotropic effects from the mutation because proinflammatory and antimicrobial activities were partitioned between A9 and the A8/A9 complex, respectively. A mutation arose in the A9 amino acid sequence and is thus present in both A9 and A8/A9 states, but we only observe effects on the A9 state. This shows that pleiotropic constraint can be reduced when protein functions are partitioned amongst different protein states.

These findings reveal the diversity of pleiotropic roles that a single mutation can play. It further shows how the deleterious pleiotropic effects of mutations can be reduced by partitioning protein functions and properties into different functional states, thus enabling the acquisition, optimization, and expansion of new protein functions. Given the vast diversity of protein functional domains and protein-protein interactions in biology, we suspect that this is a common occurrence in the evolution of protein multifunctionality.

### Conclusions

This study demonstrates the utility of placing biological findings into an evolutionary context to study the origin and mechanism of protein functions. By isolating and characterizing key intervals of evolutionary change in A9s, we have contributed to our understanding of how mammals evolved to activate and regulate inflammation and provided fresh insight into how these functions work. Our findings reveal a positive role for pleiotropy in evolution: pleiotropic substitutions can render proteins multifunctional by altering one functional state and leaving another intact, in this case leading to critical functional changes in mammalian innate immunity.

## Acknowledgements

We thank current and former members of the Harms lab for helpful discussion and input. We thank the Barber lab for use of their cell culture facility. We thank Karen Guillemin for helpful comments on the manuscript.

## Competing interests

None.

## MATERIALS AND METHODS

### Phylogenetics and ancestral sequence reconstruction

We reconstructed ancestral sequences using a previously published a phylogenetic tree of S100 proteins containing 172 sequences from 30 amniote taxa (File S2).^57^ We used PAML4 to generate maximum likelihood ancestors (marginal probability method)^87, 88^ using the previously-identified maximum likelihood (ML) substitution model (LG+Γ_8_)^89^ on the ML tree. To account for reconstruction uncertainty, we also generated “altAll” versions of each ancestor.^59^ We took every site in which the alternate reconstruction had a posterior probability > 0.20 and substituted that amino acid into the maximum-likelihood ancestor. These alternate reconstructions had an average of 12 sequence differences relative to the maximum-likelihood ancestors (Table S4). They represent a “worst case” reconstruction relative to our best, maximum likelihood reconstruction.

We also investigated the effect of topological uncertainty on our reconstructed ancestors. In the published phylogenetic analysis, A8s, A9s, A12s, and MRP126s all formed distinct and well-supported clades; however, the branching pattern between these four clades could not be resolved with high confidence.^57^ To explore how this uncertainty altered our reconstructed ancestral proteins, we constructed all 15 possible topologies for the A8, A9, A12, and MRP126 clades—i.e ((A8,A9),(A12,MRP126)), ((A8,A12),(MRP126,A9)), etc.—while maintaining species-corrected, within-clade topologies. We then optimized the tree branch lengths and substitution rates for each tree using PhyML.^90^ Finally, we used PAML to reconstruct ancA9, ancCG, and ancA8 for all 15 possible arrangements of the MRP126, A12, A8, and A9 clades. The average number of sequence differences for ancestors reconstructed using different topologies was less than or equal to the number of sequence differences between the ML and altAll reconstructions (Table S1). Further, the sites that differed were a subset of those that differed between the ML and altAll reconstructions. Thus, the altAll reconstructions account for sequence changes due to both uncertainty given the ML tree and uncertainty due to topological uncertainty.

### Cloning and mutagenesis

All S100 genes in this study were purchased as synthetic constructs in pUC57 vectors from Genscript. S100 genes (A8s, A9s, A12s, MRP126s, and ancestrally reconstructed genes) were sub-cloned into a pETDuet-1 (pD) vector (Millipore). A8s, A12s, MRP126s, and ancCGs were cloned into multiple cloning site #1 (MCS1) of the pD vector, while A9s were cloned into MCS2. For expression and purification of A8/A9 heterocomplexes (A8/A9s), pD plasmids containing an A8 gene in MCS1 and an A9 gene in MCS2 were used as previously described.^91^ Opossum A8 was sub-cloned into an MBP-LIC vector to yield a His-MBP-TEV-opA8 construct. For opossum A8/A9, the entire His-MBP-TEV-A8 construct was then sub-cloned into MCS1 of a pD vector containing a marsupial A9 in MCS2. Other S100s (A1, A5, A7, A11, A14, and P) were previously cloned into a pET28/30 vector to yield a TEV-cleavable N-terminal His tag.^67^ Cysteine-free versions of all S100 genes, as well as point mutants, were prepared using site-directed mutagenesis (Agilent).

### Protein expression and purification

Recombinant protein overexpression was conducted in *E. coli* BL21 (DE3) pLysS Rosetta cells. Cultures were innoculated in luria broth overnight at 37°C, shaking at 250 rpm, in the presence of ampicillin and chloramphenicol. The following day, 10 ml of saturated culture was diluted into 1.5 L of media with antibiotics, grown to OD_600_ = 0.6 – 1, and then induced overnight at 16°C using 1 mM IPTG. Cells were pelleted at 3,000 rpm for 20 min and stored at −20°C for no more than three months.

Lysates were prepared by vortexing pellets (3-5 g) in tris buffer (25 mM tris, 100 mM NaCl, pH 7.4) and incubating for 20 min at RT with DNAse I and lysozyme (ThermoFisher Scientific). Lysates were sonicated and cell debris was pelleted by centrifugation at 15,000 rpm at 4°C for > 20 min. All proteins were purified on an Äkta PrimePlus FPLC using various 5 ml HiTrap columns (HisTrap FF (Ni-affinity), Q HP (anion exchange), SP FF (cation exchange), and MBPTrap HP (MBP) - GE Health Science). A1, A5, A7, A11, A14, and S100P were purified using a a TEV-cleavable His tag strategy used by our lab previously^1, 3, 4^. All other S100s, except for opossum A8 and opossum A8/A9, were purified in three steps using Ni-affinity chromatography in the presence of calcium followed by two rounds of anion exchange chromatography at different pHs. For Ni-affinity chromatography, proteins were eluted over a 50 ml gradient from 25-1000 mM imidazole in tris buffer. Peak elution fractions were pooled and placed in dialysis overnight at 4°C in 4 L of tris buffer (calcium-free) adjusted to pH 8. Anion exchange chromatography was then performed the following day over a 50 ml gradient from 100-1000 mM NaCl in pH 8 tris buffer. Fractions containing majority S100 were pooled and analyzed for purity on an SDS-PAGE gel. If trace contaminants remained, an additional anion exchange step was performed at pH 6 using the same elution strategy as for the previous anion exchange step.

Opossum A8 and A8/A9 lysates were prepared as above and then flowed over a nickel column, eluting over a 50 ml gradient from 25-1000 mM imidazole in tris buffer. Peak elution was pooled and the MBP tag was cleaved by incubation with ∼1:5 TEV protease at 4°C overnight in 4 L of tris buffer. The MBP tag was then removed by flowing the sample over an MBPTrap column, step-eluting with 10 mM maltose. Additional MBP columns were run until all MBP was removed from the purified protein, assessed by SDS-PAGE. If necessary, an additional anion exchange step at pH 8 was performed to complete purification. All purified proteins were dialyzed overnight at 4°C in tris buffer + 2 g/L Chelex-100 resin (Biorad), flash-frozen the following day in liquid nitrogen, and stored at −80°C.

### Biophysical and biochemical characterization

For all experiments, protein aliquots were thawed fresh from freezer stocks and were either dialyzed in the appropriate experimental buffer overnight at 4°C or exchanged 3X into experimental buffer using 3K microsep spin concentrator columns (Pall Corporation). All samples were filter-sterilized using 0.1 um spin filters (EMD Millipore) prior to measuring concentration and using in experiments. Thawed aliquots were used for no more than one week before discarding. All concentrations were measured by Bradford assay and correspond to micromolar dimeric protein.

For *in vitro* proteolytic susceptibility experiments, proteins were dialyzed or exchanged into tris buffer + 1 mM CaCl_2_. 12.5 uM S100 protein was treated with 5 uM monomeric Proteinase K from *Tritirachium album* (Sigma Aldrich) in thin-walled PCR tubes, which were held at a constant temperature of 25°C over the course of the experiment using a thermal cycler. Proteinase K activity was quenched at different time points by directly pipetting an aliquot of the reaction into an equal volume of 95% Laemmli SDS-PAGE loading buffer + 5% BME at 95°C in a separate thermal cycler. Time points were analyzed via SDS-PAGE, and gels were quantified by densitometry using in-house gel analysis software (https://github.com/jharman25/gelquant). An exponential decay function (*A*_*O*_*e^-kt^*) was fit to the data to extract the decay rate, floating *A_O_* and *k*.

Oligomeric states were measured using a superose 12 10/300 GL size exclusion column (Amersham Biosciences) with in-line concentration detection using refractive index (RI) and particle mass measured using a multiangle laser light scattering (MALS) instrument (Dawn Heleos, Wyatt Technology). Samples were concentrated to 0.5-2 mg/ml in tris buffer + 0.5 mM CaCl_2_, 0.1 um sterile-filtered, and analyzed at a flow rate of 0.2 ml/min. Data were processed using manufacturer’s software (Astra).

Circular dichroism (CD) and chemical denaturation experiments were performed using a Jasco J-815 CD spectrometer and spectroscopy-grade guanidine hydrochloride (gdn-HCl). Chemical denaturation was performed using 25 uM dimeric protein in tris buffer with CaCl_2_, with tris substituted for spectroscopy-grade trizma. Reversible unfolding and refolding curves were constructed by making concentrated 100 uM protein stocks in either buffer or 6M gdm and then preparing protein dilutions in various concentrations of gdn-HCl in buffer. Samples were left to equilibrate in denaturant for a minimum of three hours to allow for equilibration and were then analyzed by CD. Unfolding/refolding equilibration was confirmed by comparing unfolded vs. refolded protein at the same concentration. CD signal was quantified at 222 nm in a 1 mm cuvette using a 1 nm bandwidth, standard sensitivity, and 2 second D.I.T. HT voltage was < 600 V for all measurements. Apparent unfolding kinetics studies were performed using the above conditions by spiking concentrated protein stock directly into 6M gdm and immediately monitoring CD signal at 222 nm.

### Functional Assays

The antimicrobial activity of S100s was measured against *S. epidermidis* using a well-established assay.^32, 36, 40, 41, 92^ The day before, a 5 ml starter culture of *S. epidermidis* in tryptic soy broth (TSB) was grown overnight. The next day, the culture was diluted ∼1:100 in TSB and grown for approximately 2 hours to an OD600 of ∼0.8. Immediately prior to experiment, the *S. epidermidis* culture was again diluted 1:100 at a ratio of 62:38 experimental buffer (25 mM tris, 100 mM NaCl, 3 mM CaCl_2_, pH 7.4):TSB. S100 proteins were exchanged into experimental buffer. Each well of a sterile 96-well plate was prepared with 40 ul of *S. epidermidis* diluted in experimental buffer + TSB, S100 protein at the desired concentration in experimental buffer, and then filled to 200 ul, maintaining a ratio of 62:38 experimental buffer:TB. *S. epidermidis* growth was monitored on a plate reader, measuring OD600 every 15 minutes for 13 hours. Each measurement was collected in technical triplicate and background-subtracted using a blank containing experimental buffer and TSB alone. Protein samples were confirmed to lack bacterial contamination by measuring S100 protein growth in experimental buffer and TSB lacking *S. epidermidis*.

All plasmids, cell culture conditions, and transfections for measuring the activity of S100s against TLR4s were identical to those previously described.^38,39, 57, 60^ Lipopolysaccharide *E. coli* K-12 LPS (LPS - tlrl-eklps, Invivogen) aliquots were prepared at 5 mg/ml in endotoxin-free water and stored at −20°C. Working solutions were prepared at 10 ug/ml and stored at 4°C to avoid freeze-thaw cycles. S100 proteins were prepared by exchanging into endotoxin-free PBS and incubating with an endotoxin removal column (Thermo Fisher Scientific) for 2 hours. S100 LPS contamination was assessed by measuring activity with and without Polymixin B, an LPS chelating agent (Figure S6). LPS (200 ng per 100 ul well) or S100 (0.8, 0.4, 2, 4, or 5 uM dimer) treatments were prepared by diluting in 25:75 endotoxin-free PBS:serum-free Dulbecco’s Modified Eagle Media (DMEM – Thermo Fisher Scientific). Polymixin B (PB, 200 ug per 100 ul well) was added to all S100 experimental samples to limit background endotoxin contamination activity from recombinant protein preps. Cells were incubated with treatments for 3 hours prior to assaying activity. The Dual-Glo Luciferase Assay System (Promega) was used to assay Firefly and Renilla luciferase activity of individual wells. Each NF-κB induction value shown represents the Firefly luciferase activity divided by the Renilla luciferase activity, background-subtracted using the LPS + PB activity for each TLR4 species and normalized to the activity of LPS alone for each TLR4 species to normalize between plates. All measurements were performed using three technical replicates per plate, a minimum of three biological replicates total, and a minimum of two separate protein preps.

For TLR4 activation measurements by A9 proteolytic products, 12.5 uM hA9 or hA9 M63F were incubated with 2.5 mg/ml Proteinase K immobilized to agarose at 37°C for increasing amounts of time. The reaction was quenched by spin-filtering the sample to remove Proteinase K. 2 uM A9 proteolysis treatments were then added to cells as outlined above. Western blots were performed by running an SDS-PAGE gel and transferring to a nitrocellulose membrane. Membranes were blocked using Odyssey Blocking Buffer for 1 hour, incubated with 1:1000 mouse anti-S100A9 primary antibody (M13 clone 1CD22, Abnova) for 1 hour, and incubated with 1:10,000 IRDye Goat anti-mouse 800CW IgG (H+L, Licor) for 1 hour, with 3×5 min TBST washes in between each step. Blots were imaged using the Licor Odyssey Fc imaging system.

All species cartoons were taken from the following websites: http://phylopic.org/image/c089caae-43ef-4e4e-bf26-973dd4cb65c5/, http://phylopic.org/image/aff847b0-ecbd-4d41-98ce-665921a6d96e/, http://phylopic.org/image/0f6af3d8-49d2-4d75-8edf-08598387afde/http://phylopic.org/image/dde4f926-c04c-47ef-a337-927ceb36e7ef/. We acknowledge Sarah Werning and David Liao as authors of the opossum and mouse cartoons respectively, which were made publicly available through the creative commons attributions 3.0 unported license (https://creativecommons.org/licenses/by/3.0/).

## SUPPLEMENTAL MATERIAL

**Figure S1.**
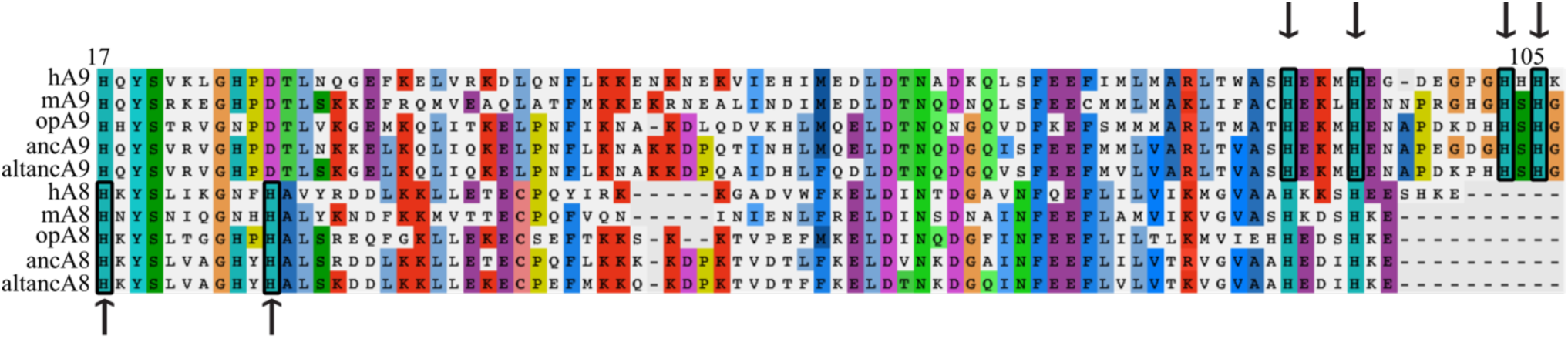
The residues composing the A8/A9 hexastidine site are conserved across mammalian and ancestrally reconstructed A8s and A9s. Human (h), mouse (m), opossum (op), maximum likelihood therian mammalian ancestors (anc), and AltAll ancestors (altanc) shown. Alignment truncated to show conservation of key hexahistidine site metal binding residues (boxed + arrows). A8s conserve two (positions 17 and 27 in human A8), while A9s conserve four (positions 91, 95, 103, and 105 of human A9). Consensus residues for alignment are highlighted.

**Figure S2.**
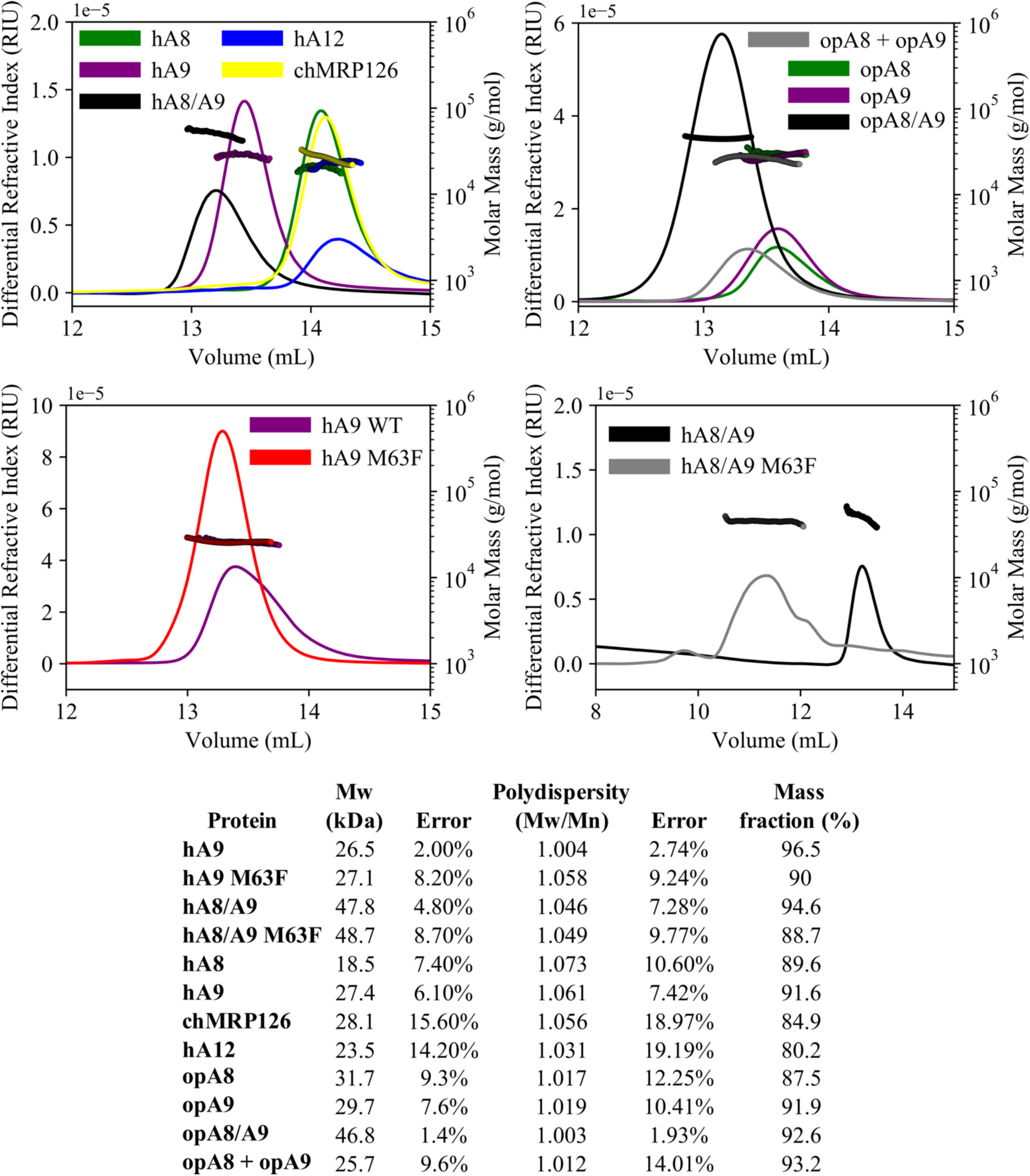
Analysis of protein oligomeric state using SECMALS. Differential refractive index (left y-axis, lines) and calculated molecular weights from light scattering detectors (right y-axis, points) for modern S100 proteins used in this study. h = human, op = opossum, ch = chicken species. Opossum A8 + A9 sample is an equimolar mixture of opossum A8 and A9 homodimers. Table below shows summary data calculated using Wyatt Astra software.

**Figure S3.**
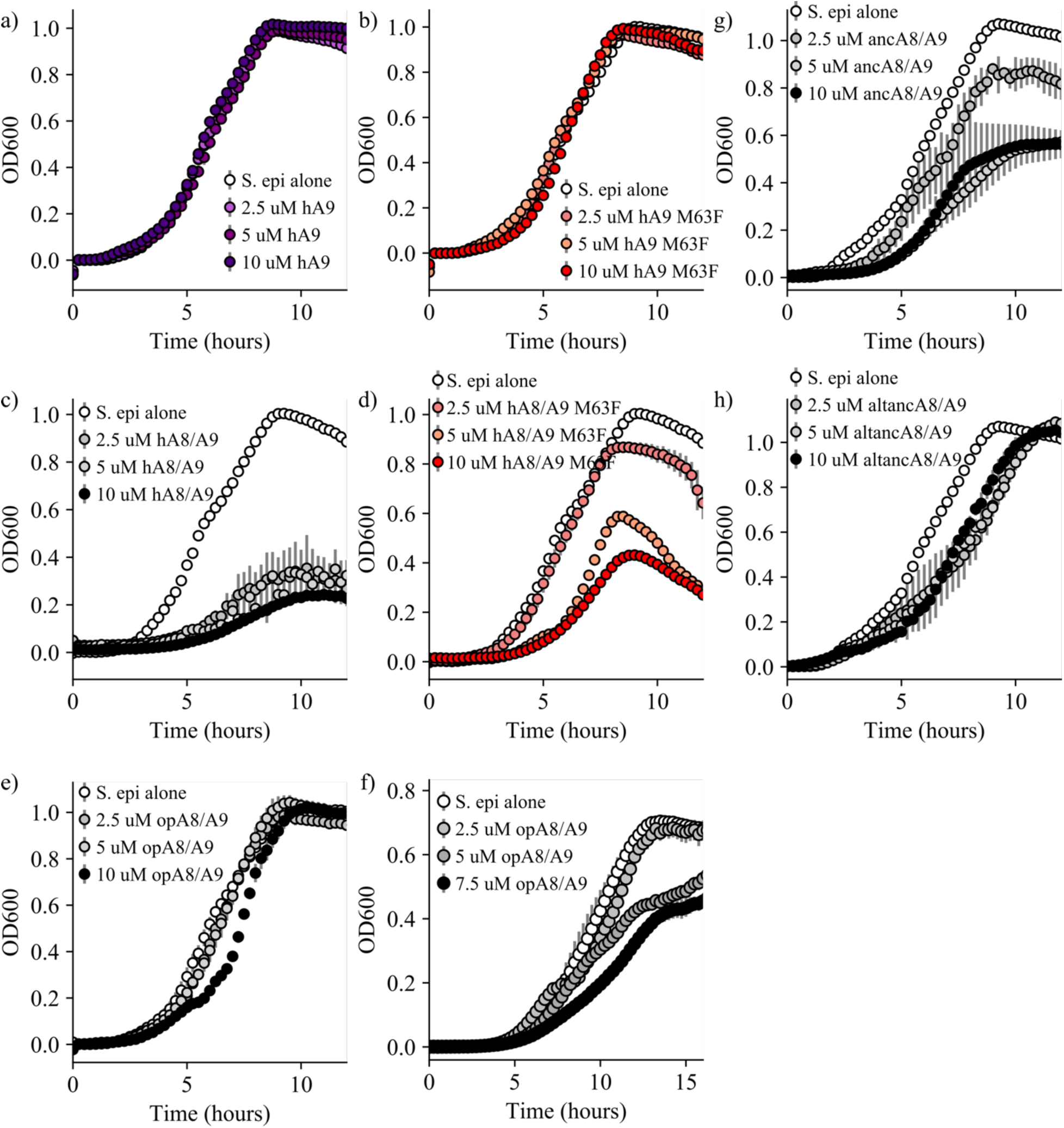
Bacterial growth in the presence of S100 proteins. Representative *S. epidermidis* growth curves in the presence of a) human A9, b) human A9 M63F, c) human A8/A9, d) human A8/A9 M63F, e) opossum A8/A9 (cysteine-free), f) opossum A8/A9 (containing cysteines), g) ancA8/A9, and h) altancA8/A9. Error bars are the standard deviation for 3 technical replicates, points show one representative biological replicate for each protein at four different concentrations.

**File S1.**
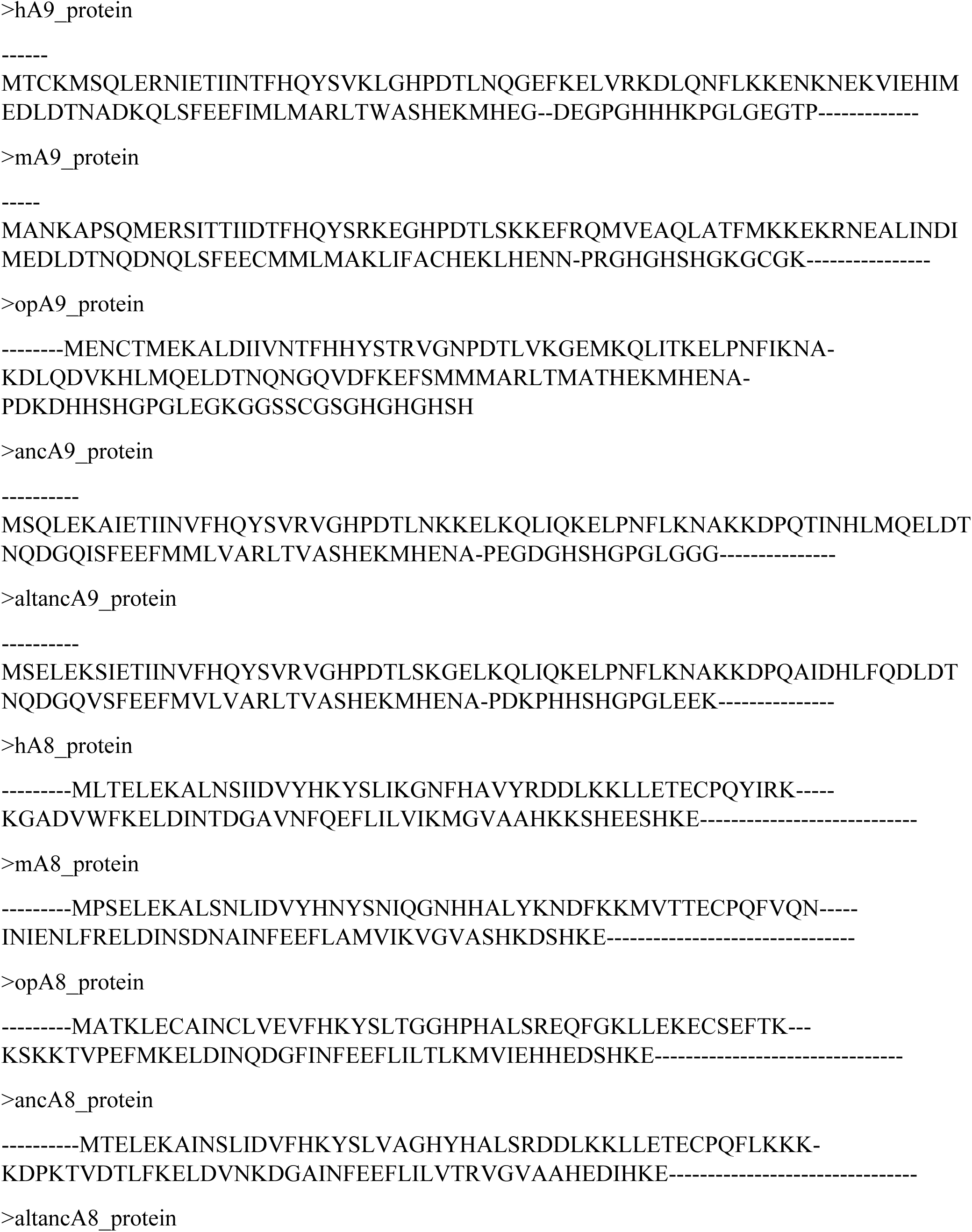
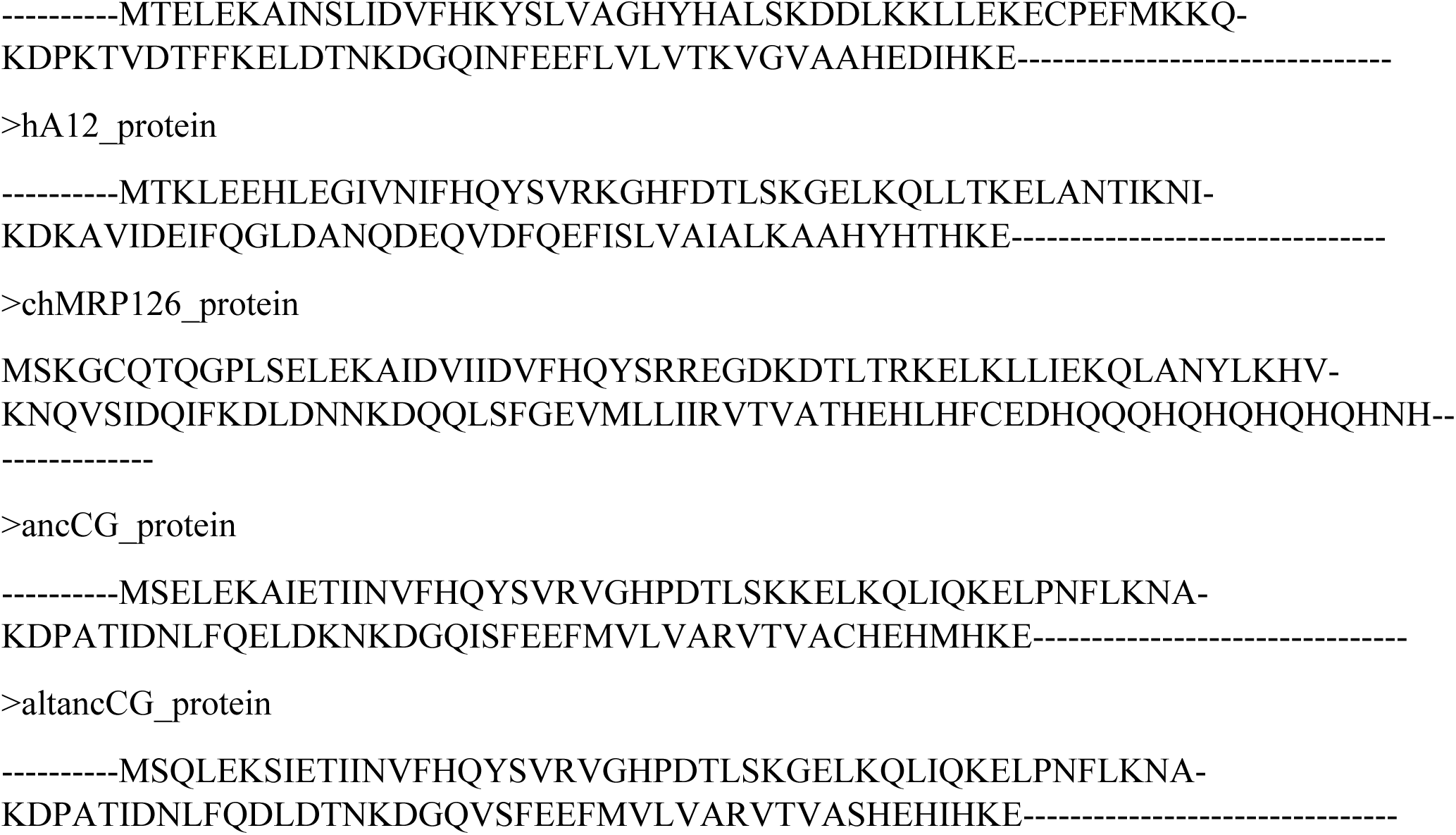
Alignment of modern and ancestrally reconstructed S100 proteins used in this study.

**Table S1.** Statistics for ancestrally reconstructed proteins.

| **Ancestrally**<br>**reconstructed protein** | **Average**<br>**posterior**<br>**probability** | **# of sites that differ**<br>**from the ML**<br>**sequence** | **Average # sites that**<br>**differ from ML**<br>**sequence for**<br>**reconstructions on**<br>**alternate topologies** |
| --- | --- | --- | --- |
| ancA9 | 0.83 | -- | 5 |
| altancA9 | 0.81 | 10 | -- |
| ancA8 | 0.88 | -- | 1 |
| altancA8 | 0.86 | 17 | -- |
| ancCG | 0.86 | -- | 8 |
| altancCG | 0.84 | 8 | -- |

**File S2:**
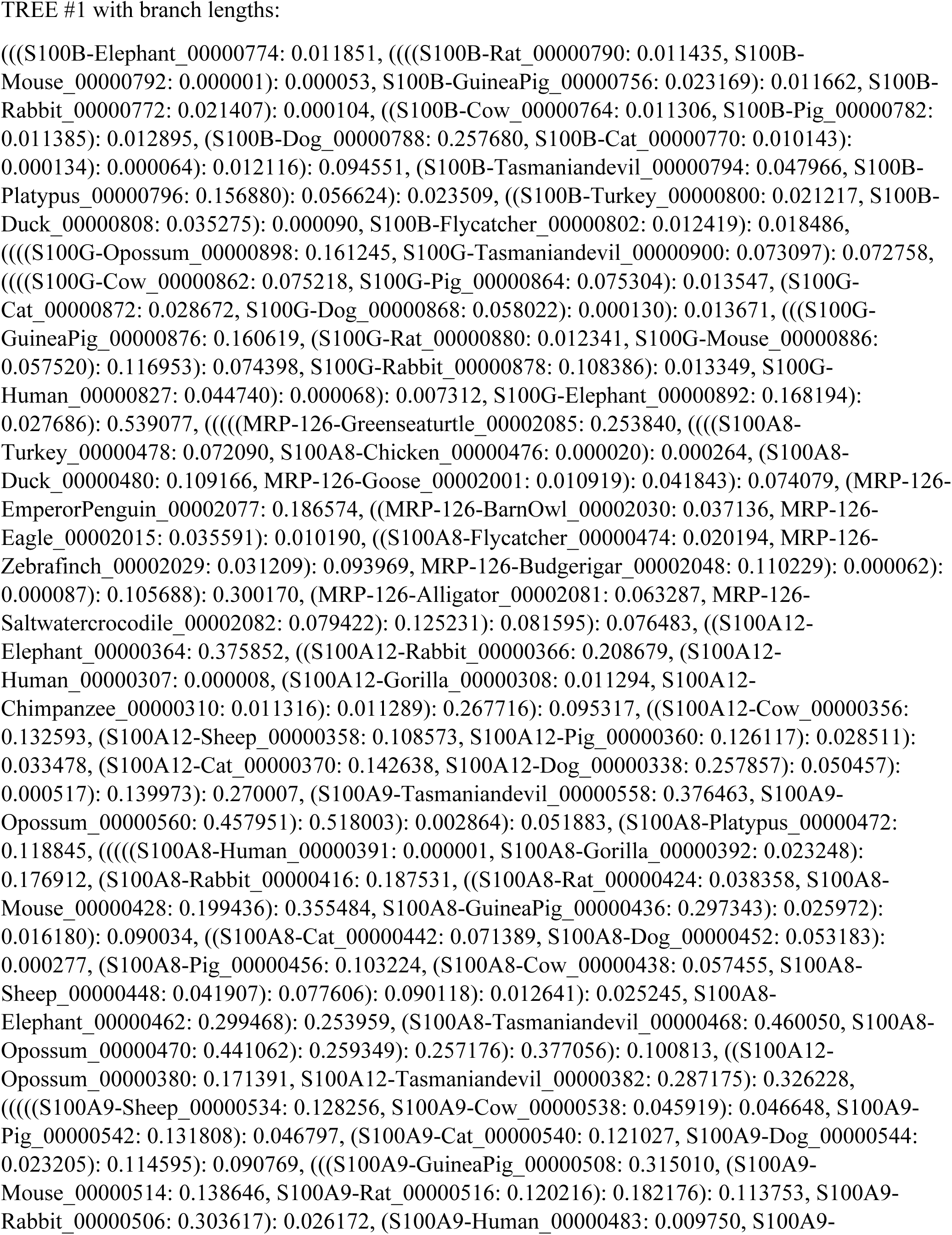
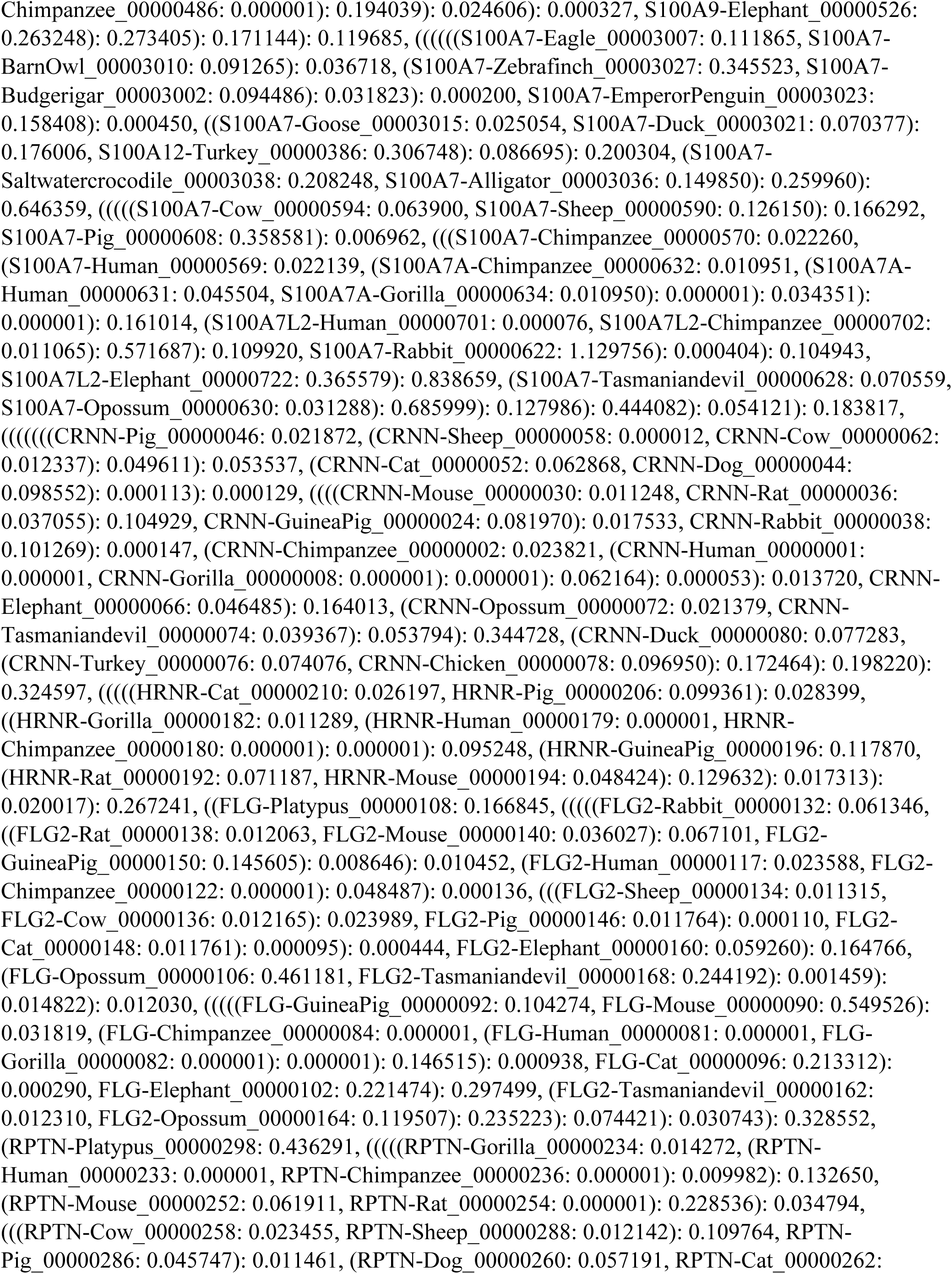
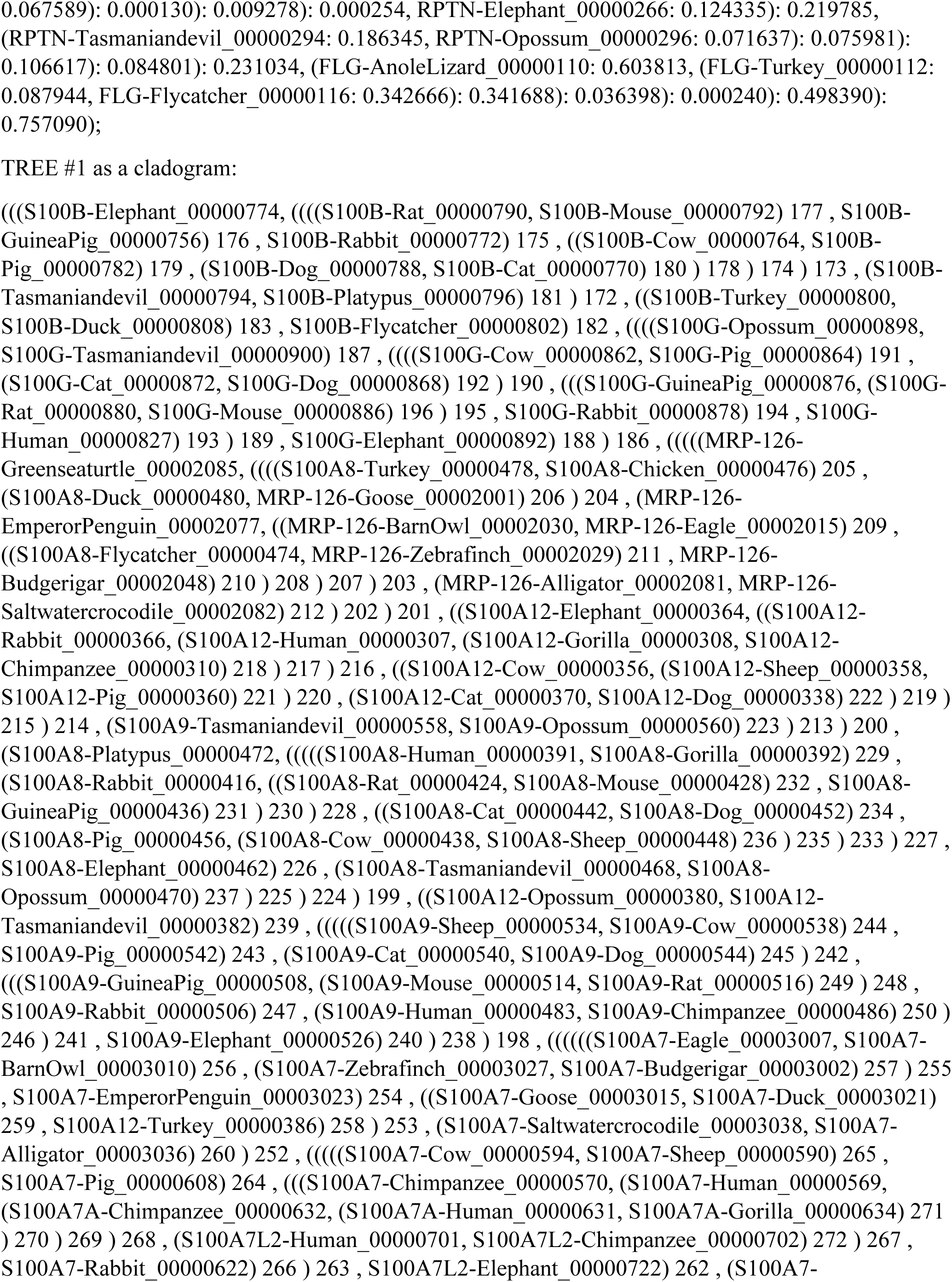
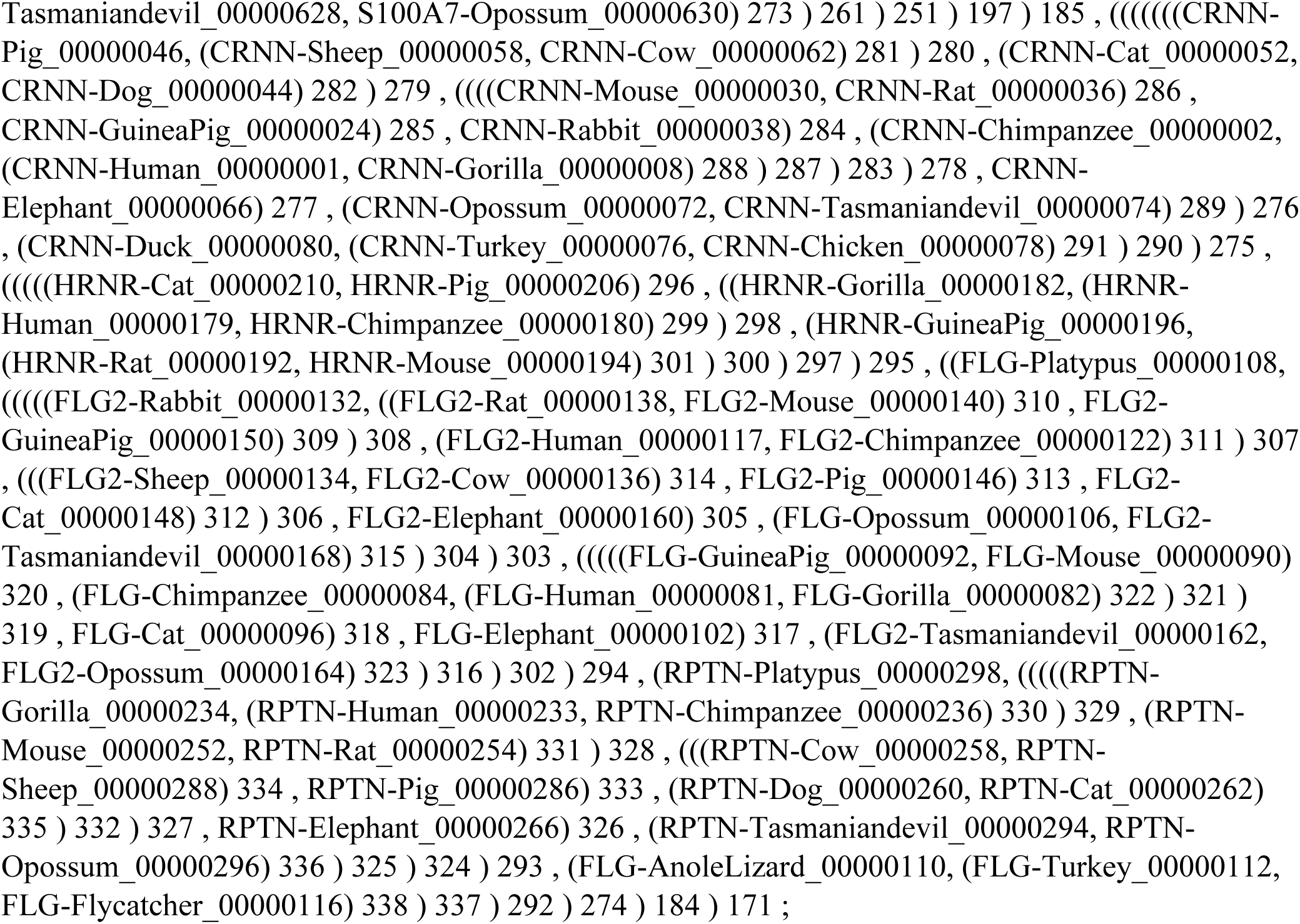
Species-corrected S100 tree used for ancestral sequence reconstruction.

**Figure S5.**
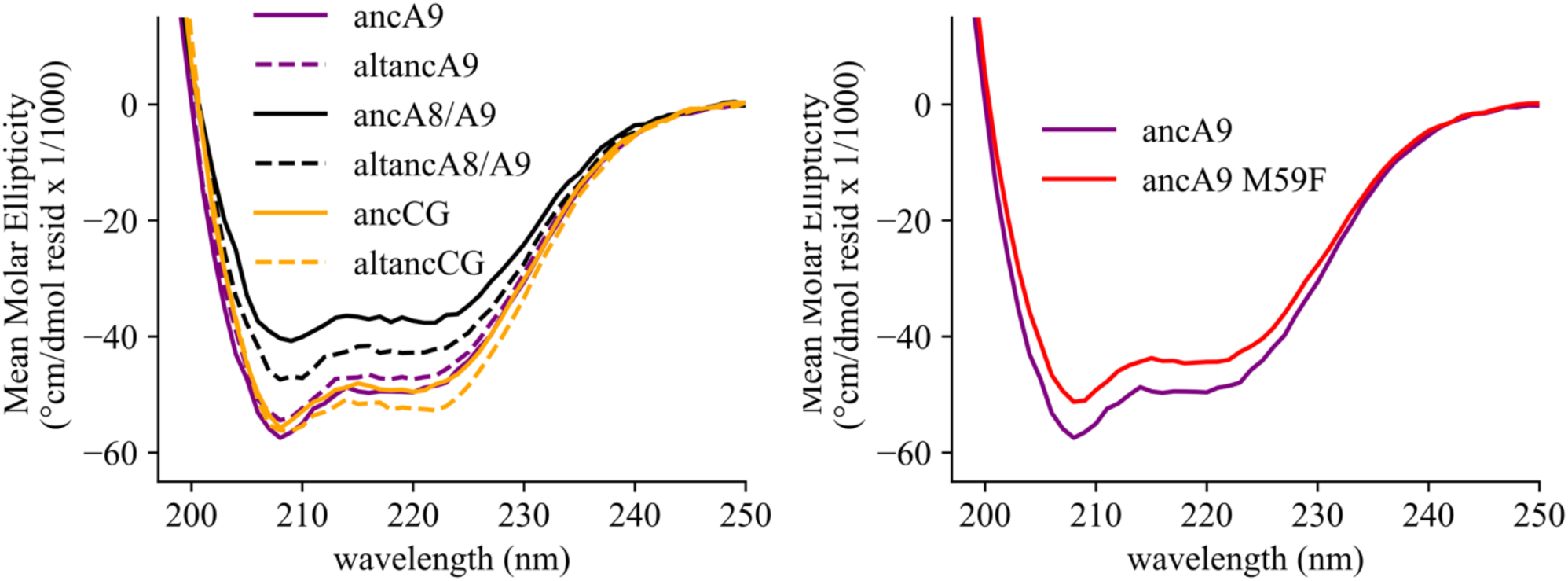
Secondary structure characterization of ancestrally reconstructed S100s by CD spectroscopy. Data shown are the average of 3 scans. Solid lines are maximum likelihood ancestral proteins, dotted lines are alt-all ancestors (colored the same as matching maximum likelihood ancestor for comparison).

**Figure S6.**
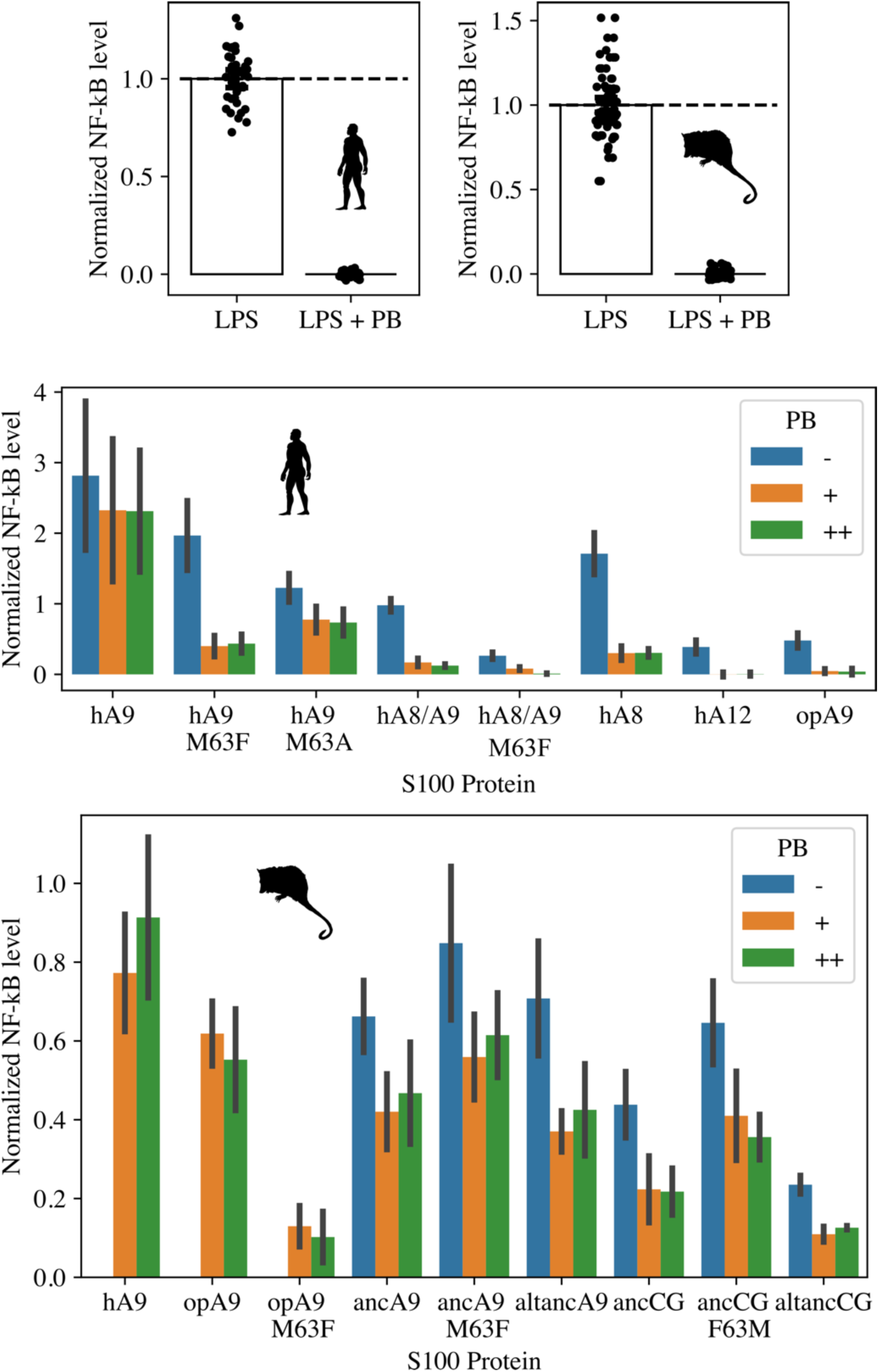
Analysis of S100 protein LPS contamination. Top panel: Activity of LPS (0.2 ng/ul) against human (left) and opossum TLR4 (right) is inhibited by the addition of Polymixin B (PB, 0.2 ug/ul). Middle and bottom panels: S100 activation of human (middle) and opossum (bottom) TLR4 with no PB, 0.2 ug/ul PB (+) and 0.25 ug/ul PB (++). No-PB data were not collected for hA9, opA9, and opA9 M60F against opossum TLR4 (bottom panel). Data were background-subtracted using the LPS + PB control and normalized to LPS activity against either human or opossum TLR4. Error bars are standard deviation.

**Figure S7.**
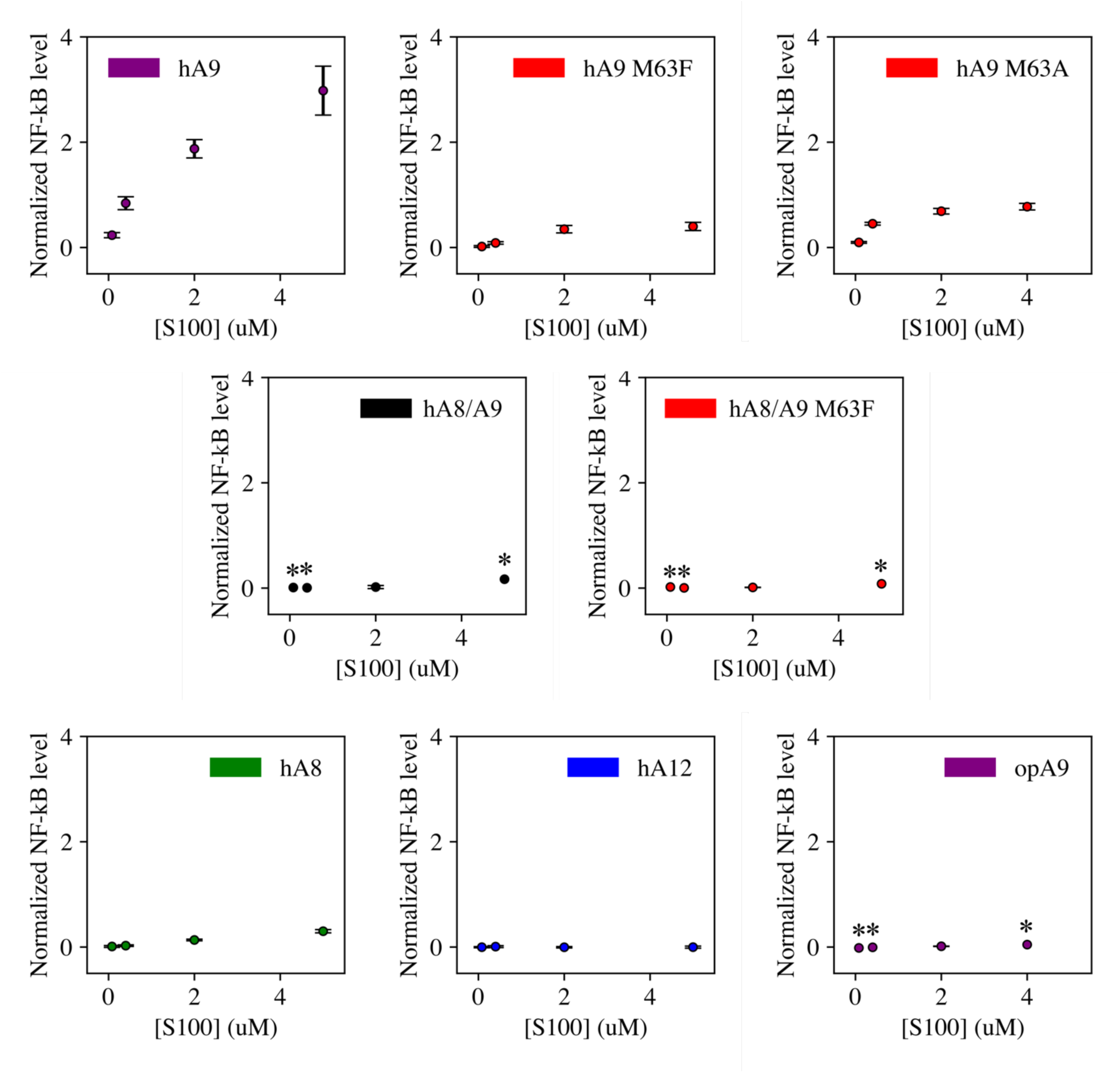
Dose curves for S100 activation of human TLR4. Points are the average of >3 biological replicates each consisting of 3 technical triplicates, error bars are standard error of the mean. An asterisk (*) indicates a concentration at which a single biological replicate was measured. Data were background-subtracted using the LPS + PB control and normalized to LPS activity against human TLR4.

**Figure S8.**
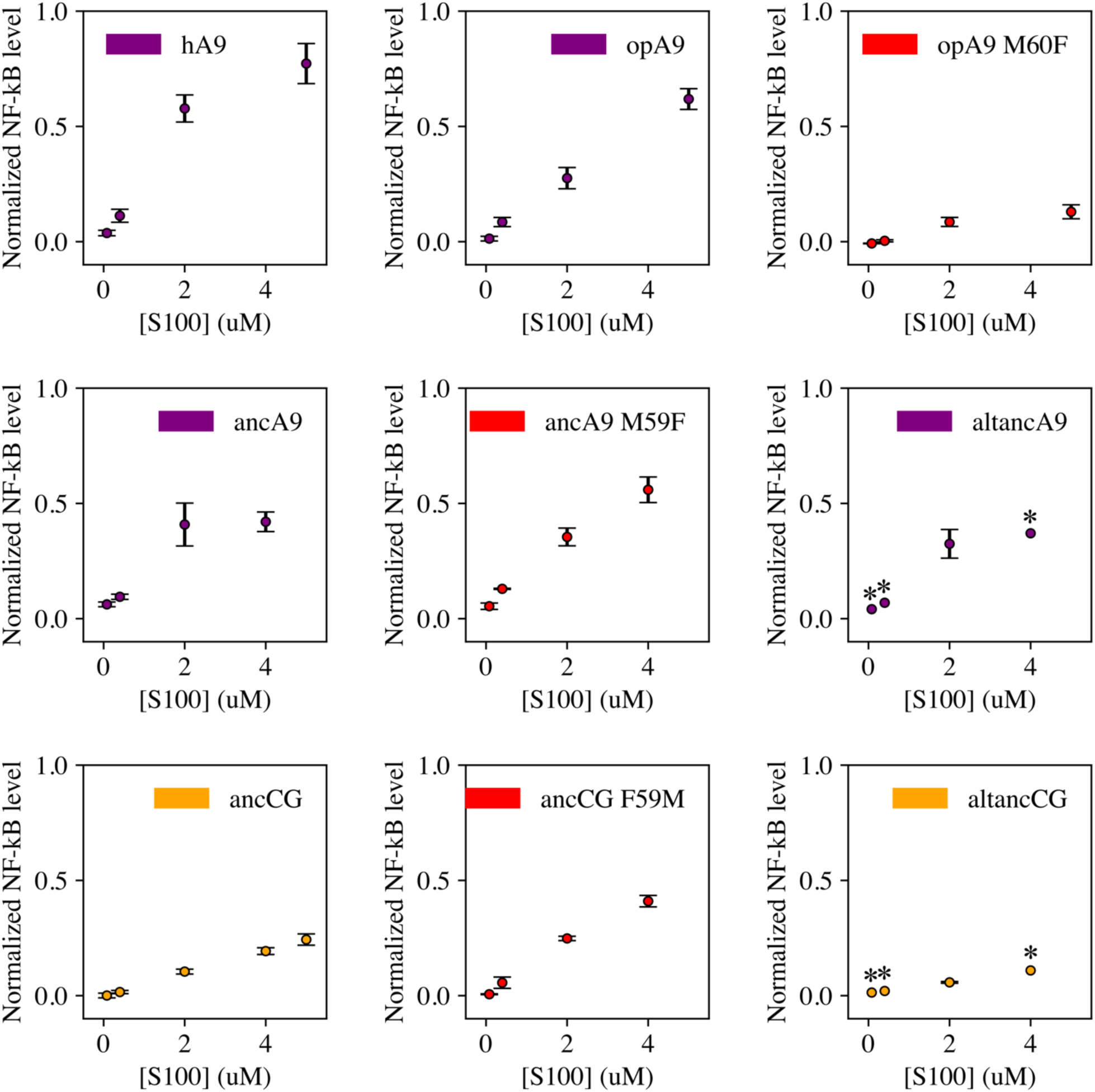
Dose curves for S100 activation of opossum TLR4. Points are the average of >3 biological replicates each consisting of 3 technical triplicates, error bars are standard error of the mean. An asterisk (*) indicates a concentration at which a single biological replicate was measured. Data were background-subtracted using the LPS + PB control and normalized to LPS activity against opossum TLR4.

**Figure S9.**
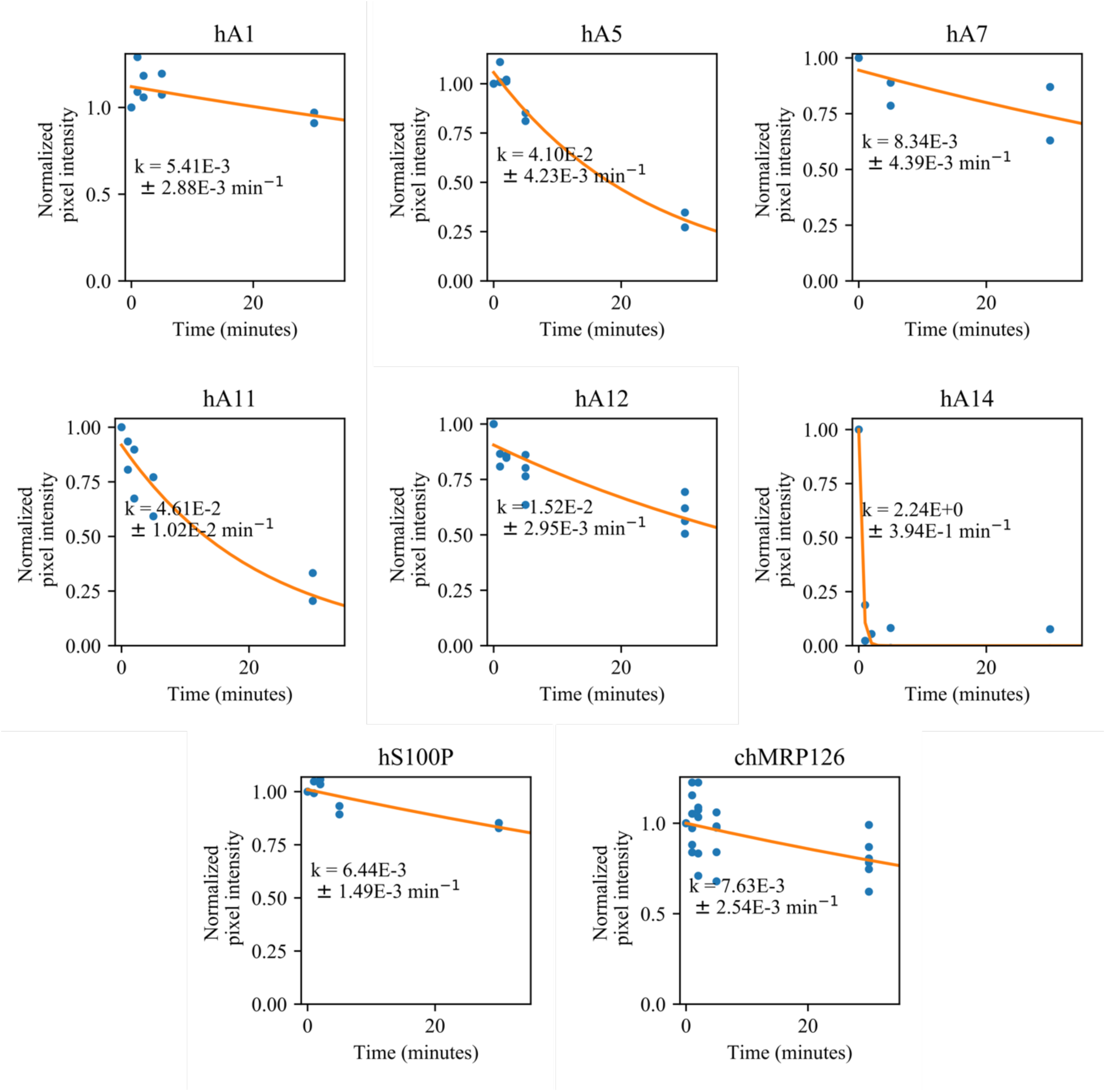
Survey of proteolytic susceptibility across modern S100 proteins. Blue dots are biological replicates, orange line is a single exponential decay fit (see methods). Protein is listed at the top. Pixel intensity was quantified by densitometry from SDS-PAGE gels.

**Figure S10.**
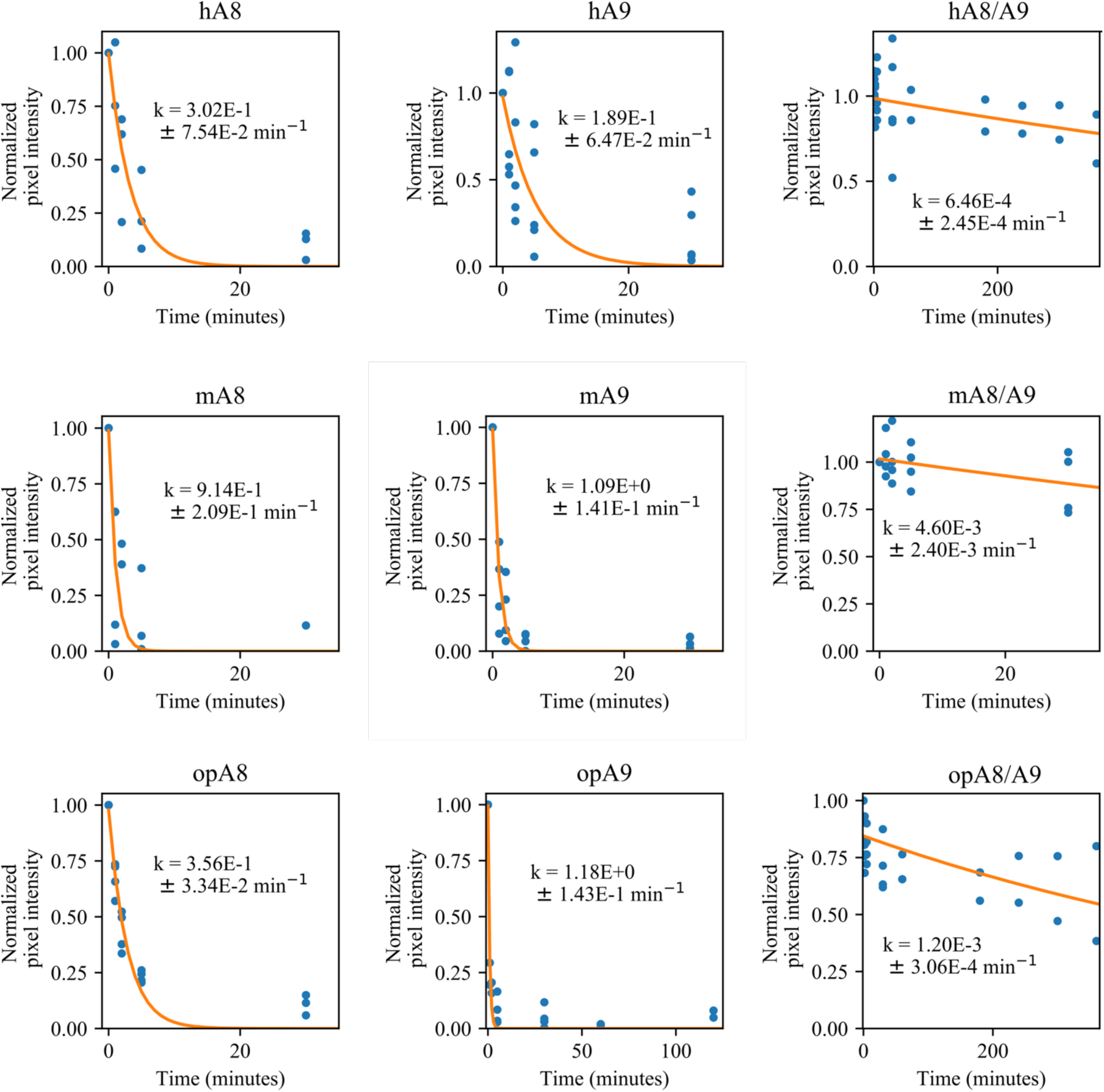
Comparison of proteolytic susceptibility for A8s, A9s, and A8/A9 complexes across mammals. Blue dots are biological replicates, orange line is a single exponential decay fit (see methods). Protein is listed at the top. Pixel intensity was quantified by densitometry from SDS-PAGE gels. Longer time points were collected for proteins with slower degradation rates (see x-axis).

**Figure S11.**
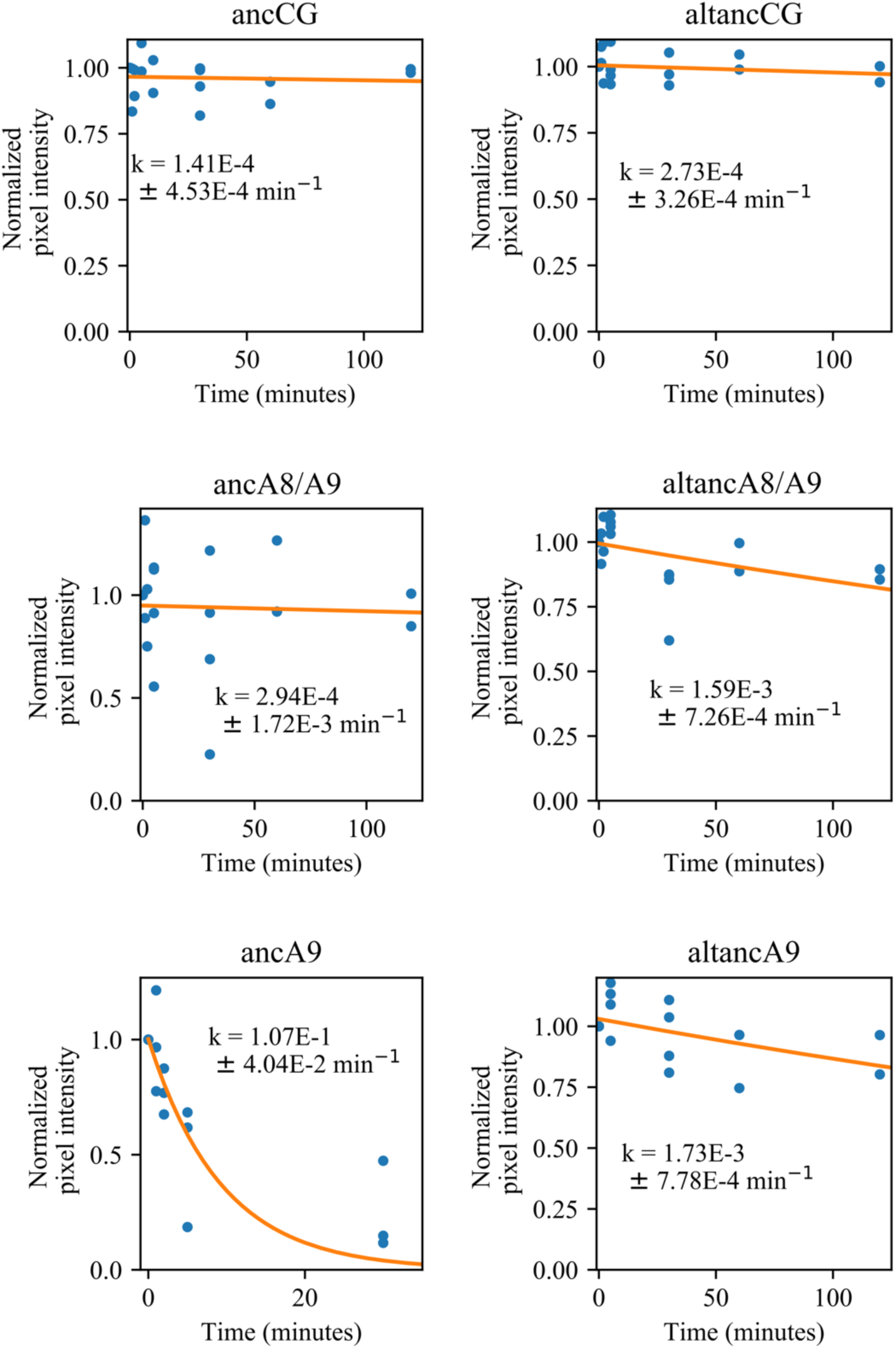
Comparison of proteolytic susceptibility for ancestrally reconstructed S100s. Blue dots are biological replicates, orange line is a single exponential decay fit (see methods). Protein is listed at the top. Pixel intensity was quantified by densitometry from SDS-PAGE gels. Longer time points were collected for proteins with slower degradation rates (see x-axis).

**Figure S12.**
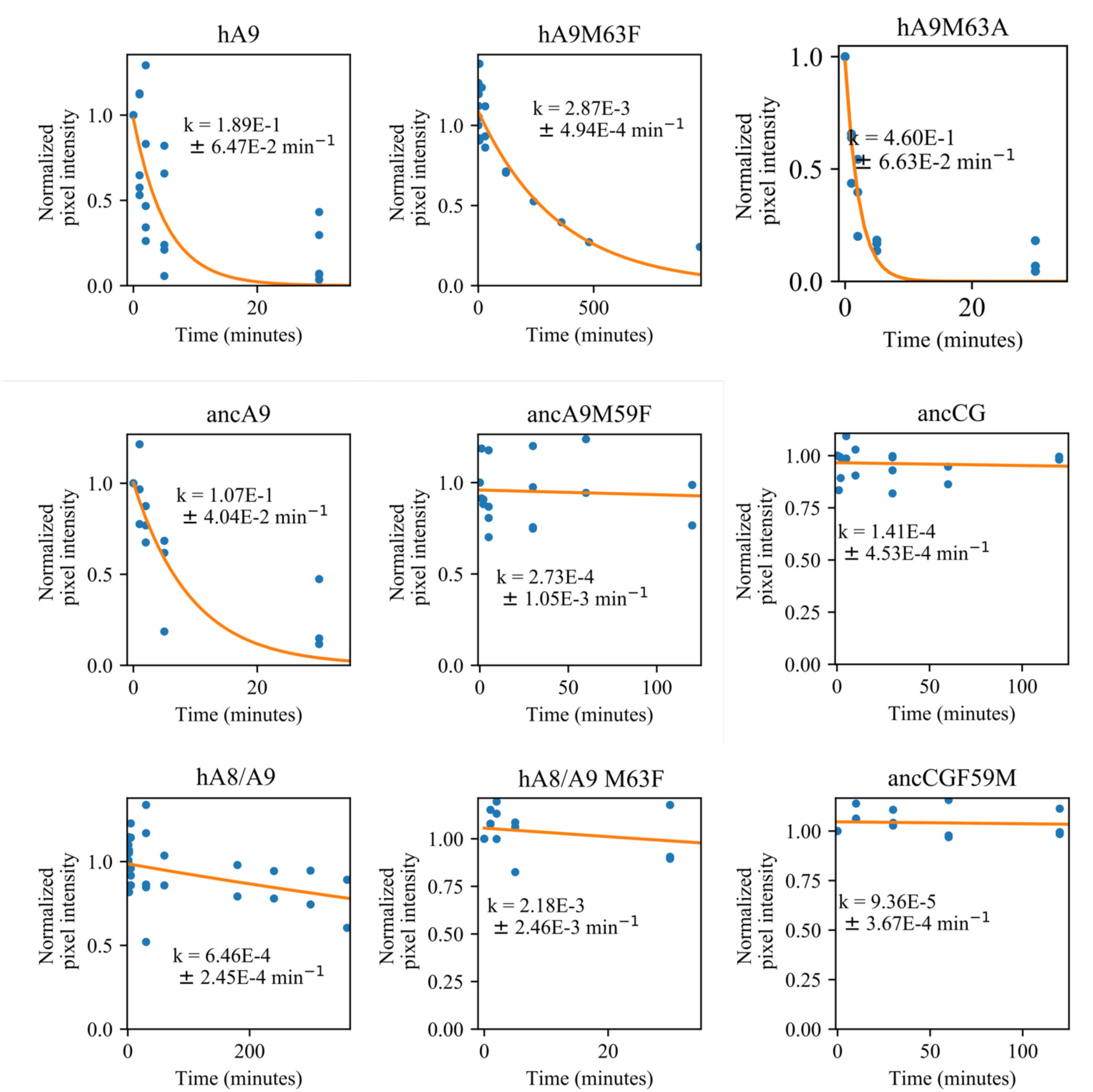
Comparison of proteolytic S100 protein mutants at position 63. Blue dots are biological replicates, orange line is a single exponential decay fit (see methods). Protein is listed at the top. Pixel intensity was quantified by densitometry from SDS-PAGE gels. Longer time points were collected for proteins with slower degradation rates (see x-axis).

**Figure S13.**
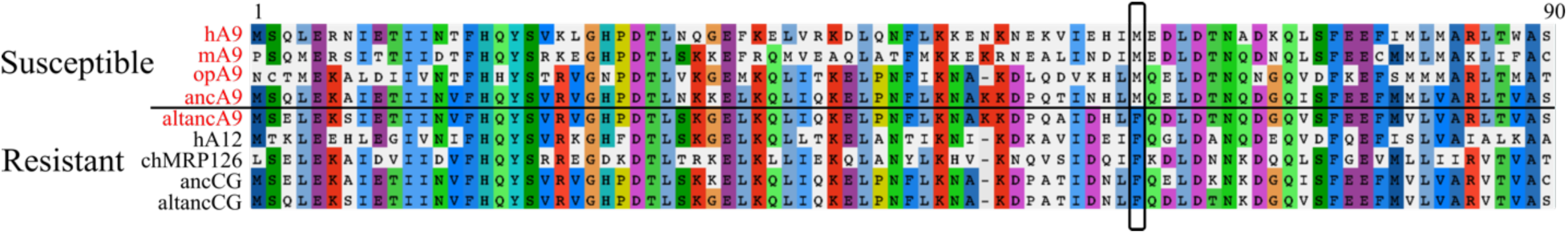
Changes at position 63 correlate with A9 proinflammatory activity and proteolytic resistance. S100 protein sequences are grouped into proteolytically susceptible (top) or resistant and potently proinflammatory (red text) or not (black text). Only the first 90 residues out of 114 total were examined as the disordered A9 tail (residues ∼93-114) are highly variable and the tail is dispensable for A9 proinflammatory activity. Residues are colored when found to be the consensus residue for a column.

**Figure S14.**
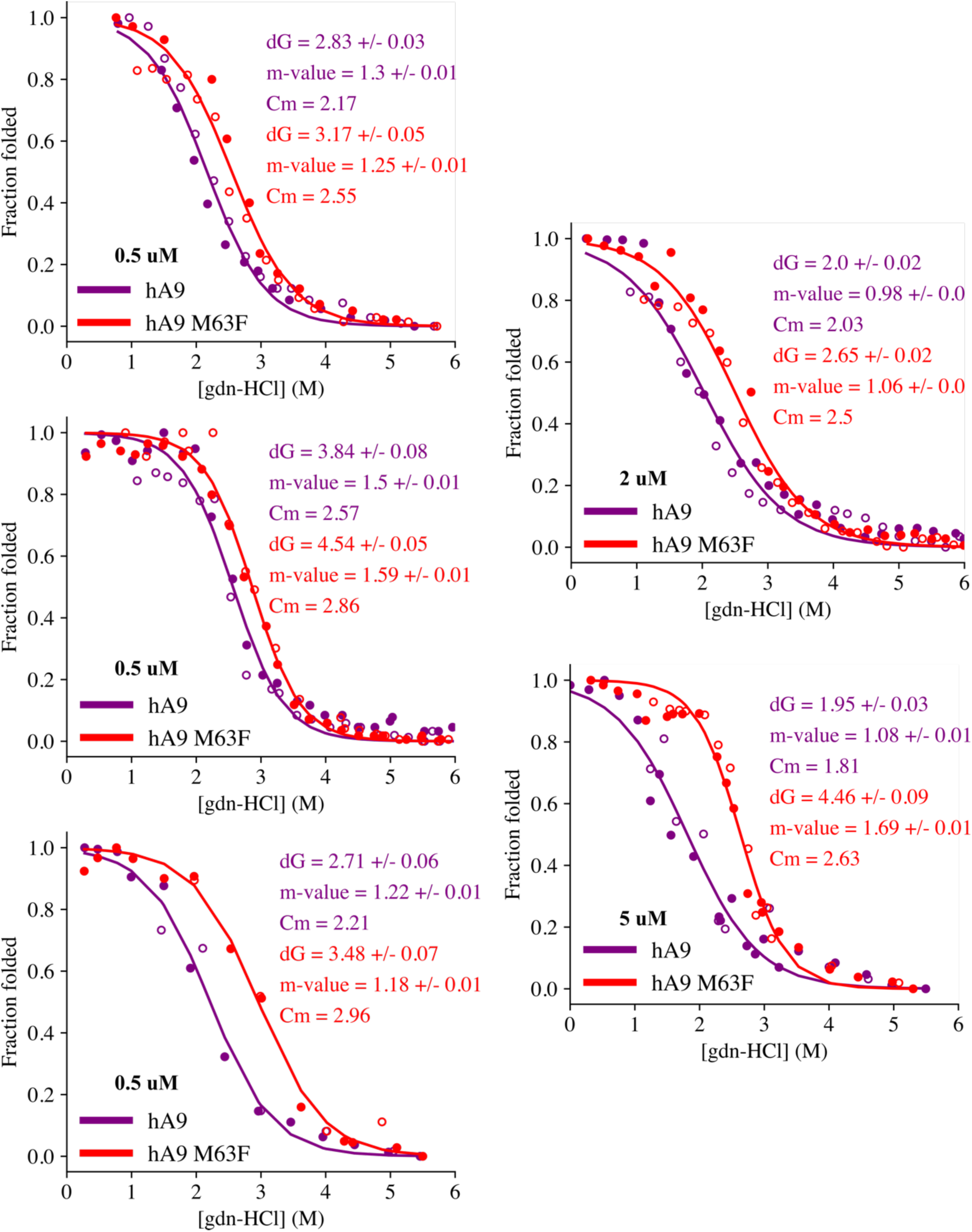
Chemical denaturation of hA9 and hA9 M63F. Each graph is a single biological replicate. Filled circles are unfolding, empty circles are refolding. A two-state unfolding model was fit to the data, shown as a solid line. Because A9 is a dimer and could therefore exhibit concentration-dependent folding properties, we verified that the C_m_ for M63F was higher than A9 at 0.5, 2.0, and 5 μM protein. Protein concentrations are noted on graphs.

**Figure S15.**
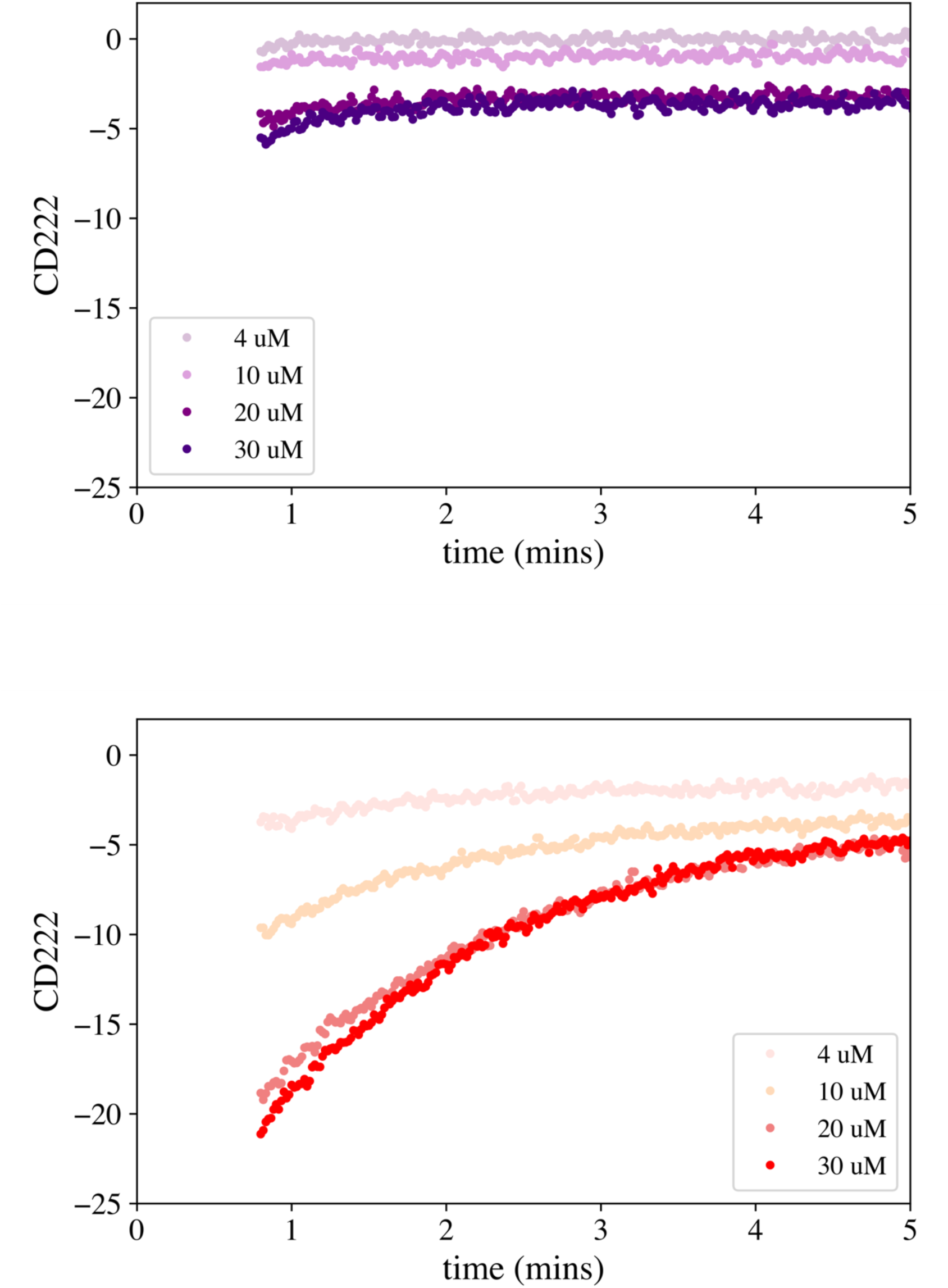
Unfolding kinetics of hA9 and hA9 M63F. Time course measurement of hA9 (top) and hA9 M63F (bottom) unfolding upon addition of 6M gdn-HCl. Each curve is a single replicate at one concentration, monitoring CD signal at 222 nm (y-axis).

## Notes

**Funding sources:** This research was funded by grants from the American Heart Association (AHA16 15BGIA22830013, MJH) and the National Institutes of Health (NIH-3R01GM117140-03S1, MJH; NIH-T32GM007413, JLH, ANL). MJH is a Pew Scholar in the Biomedical Sciences, supported by The Pew Charitable Trusts. The funders had no role in study design, data collection and analysis, decision to publish, or preparation of the manuscript.

